# Dynamic mechanical sorting biases niche occupancy and lineage outcome during development of bilayered glandular epithelia

**DOI:** 10.64898/2026.11.05.761865

**Authors:** Robin P. Journot, Tom Deshayes, Louise Harari, Alexandre Barthelemy, Vasudha Srivastava, Candice Merle, Claudia Carabana, Marisa M. Faraldo, Philip Allan Seymour, Martti Maimets, Zev J. Gartner, Jean-Léon Maître, Kim B. Jensen, Silvia Fre

## Abstract

Tissue compartmentalization into basal and luminal domains is a hallmark of glandular epithelia, yet how fate boundaries are established and stabilized during dynamic morphogenesis remains a major unresolved question. Here, using mouse embryonic explants, organoids, quantitative live imaging and genetic perturbations, we show that basal-luminal compartmentalization is not maintained by fixed boundaries, but through continuous positional corrections. We show that differences in cellular adhesion and actomyosin contractility drive these positional corrections by promoting active cell sorting. We further find that these mechanical differences influence lineage specification by controlling cell positioning rather than directly instructing cell fate. Before commitment, mosaic, but not uniform, perturbation of cell mechanics alters niche occupancy and biases lineage outcome, indicating that relative mechanical properties determine access to fate-instructive niches. Conversely, perturbation of Notch and YAP remodels compartment-specific mechanical properties and cell-positioning dynamics, revealing a reciprocal coupling between lineage state and mechanical sorting. Conservation of this mechanism across four glandular epithelia of distinct origins and functions identifies dynamic mechanical sorting as a general strategy for maintaining robust tissue compartmentalization during the extensive growth and cellular rearrangements of branching morphogenesis.

## Introduction

Building a functional tissue requires cells to acquire distinct identities, but also to occupy defined positions within the tissue. Yet morphogenesis continuously challenges the coordination between this process as cells divide, move and rearrange. Branched exocrine glands exemplify this problem; despite their diverse origins and functions, mammary, lacrimal, salivary and prostatic epithelia share a conserved bilayered organization in which basal and luminal lineages occupy the outer and inner epithelial compartments, respectively. During embryogenesis, these compartments emerge as epithelial placodes invaginate and expand into branched ductal networks while progenitor cells commit to basal and luminal fate^1–4^. How this spatial organization is established and maintained throughout branching morphogenesis remains unknown.

Several mechanisms have been proposed to explain how tissue boundaries are preserved. One framework emphasizes lineage restriction and boundary-associated signalling, whereby cells on either side of an interface acquire distinct identities while local signalling interactions and, in some contexts, extracellular matrix deposition restricts inappropriate fate transitions and heterotypic mixing^5–8^. A complementary physical framework emphasizes differential cell sorting, in which differences in adhesion and cortical tension could generate effective interfacial tensions that favor segregation between neighbouring cell populations^9–12^. In the context of exocrine gland development, cell adhesion and actomyosin contractility are established organizers of epithelial architecture. Mammary epithelial cells can self-organize into bilayers, and perturbations of adherens junctions or contractility disrupt this organization^13–15^. During embryonic salivary gland morphogenesis, spatial differences in the balance between cell-cell and cell-matrix adhesion drive epithelial remodeling and can recapitulate key morphogenetic behaviors in reconstitution assays^16^. However, whether basal-luminal segregation is maintained by restricting cell mixing or by repositioning cells after transient exchanges remains unclear, as does whether cell position influences or simply reflects lineage commitment.

Cell position is intimately linked to lineage identity, as cells occupying different epithelial domains experience distinct local signalling environments. We previously showed that Hippo/YAP, Notch and p63, conserved regulators of stemness and lineage specification across bilayered epithelia, are modulated by spatial context, indicating that cell position contributes to fate specification^17^. During morphogenesis, however, cells continuously change position, raising the question of whether lineage choice depends on transient position or sustained niche occupancy. How frequently cells exchange between basal and luminal compartments before commitment, and whether these dynamics influence fate, remains unknown.

Here, we combine quantitative live imaging of embryonic explants with organoid models and genetic perturbations to dissect basal-luminal organization during gland morphogenesis. We find that sharp compartmentalization persists despite frequent transient exchanges across the basal-luminal interface. Relative differences in adhesion and actomyosin contractility drive positional correction through active cell sorting. Before lineage commitment, mosaic, but not uniform, perturbation of cellular mechanical properties biases cell position and fate, indicating that mechanics influences lineage choice by controlling access to instructive niches. Conversely, lineage regulators remodel mechanical properties and positioning dynamics, revealing a reciprocal coupling between mechanics, position and fate that supports robust compartmentalization during branching morphogenesis. Its conservation across glandular epithelia with distinct developmental origins and functions identifies mechanical control of cell positioning as a reusable morphogenetic principle for coupling tissue organization to lineage specification.

## Results

### Basal-luminal compartmentalization is maintained as a dynamic, self-correcting steady state

To determine whether epithelial boundary maintenance relies on static positional constraints or continuous dynamic cell rearrangements, we first focused on developmental stages where basal and luminal lineages are already molecularly distinct. We established long-term live imaging of mouse embryonic mammary, lacrimal, salivary, and prostatic explants, by culturing them *ex vivo* with their endogenous mesenchyme at an air-liquid interface^17^ (Figure 1A, B; S1A; Video S1-4; Methods). In parallel, we established matched organoid cultures enabling high-resolution imaging and single-cell tracking over several days in culture (Figure 1A; S1B, C; Methods). In explants, branch architecture varied across tissues (Figure S1D, E), however, bilayer compartmentalization was conserved across tissues, establishing a common experimental setup to interrogate boundary dynamics. In both explants and organoids, cells segregated into two spatially and molecularly distinct domains; an outer basal layer, expressing the basal markers K14/p63, and an inner luminal compartment marked by cytokeratin K8 and the Notch target HES1 (Figure 1C, D; S1F, G). In fixed samples, the interface appeared sharply demarcated, consistent with a stable compartment boundary.

**Figure 1:**
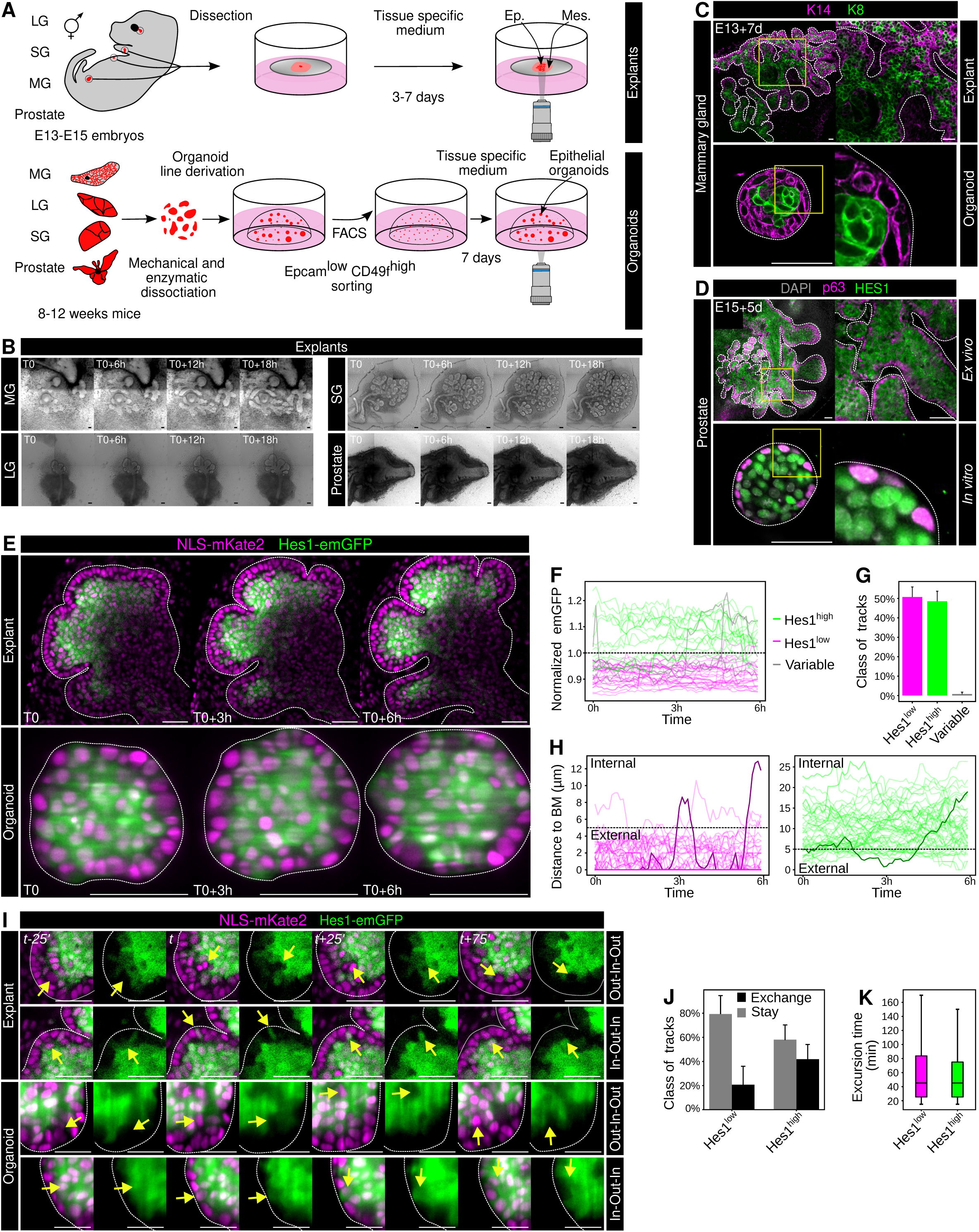
Tissue compartmentalization is maintained during branching morphogenesis of glandular epithelia despite extensive cellular rearrangements. **A:** Schematic representation of the workflow for embryonic explant culture (top) and organoid generation (bottom) from FACS-sorted adult basal cells (BCs) isolated from the mammary gland (MG), lacrimal gland (LG), salivary gland (SG), and prostate. **B:** Brightfield imaging of embryonic explants showing recapitulation of branching morphogenesis *ex vivo*. T0 corresponds to E13+5d (MG), E13+2d (SG), E15+3d (LG and Prostate). **C:** Confocal imaging of mammary embryonic explants and organoids stained for K14 (magenta) and KS (green). **D:** Confocal imaging of prostate embryonic explants and organoids stained for p63 (magenta) and HES1 (green). **E:** Optical sections from live imaging of an SG explant and an organoid expressing fluorescent reporters for nuclei (NLS-mKate2) and HES1 activity (Hes1-emGFP). **F:** Normalized Hes1-emGFP fluorescence intensity over time for individual cells within mammary organoids. One hundred representative cell tracks are shown. One representative track is highlighted per class. **G:** Quantification of Hes1-emGFP track classes: variable (CV > 0.2), Hes1^high^ (Hes1_norm_ > 1 and CV < 0.2), and Hes1^low^ (Hes1_norm_ < 1 and CV < 0.2). **H:** Absolute distance of individual cells from the basement membrane (BM) over time within organoids. One hundred representative tracks are shown, stratified by Hes1-emGFP track class. One representative track undergoing compartment exchange is highlighted by class. **I:** Optical sections from live imaging of an SG explant and an organoid illustrating cell movements between the internal and external epithelial compartments. Yellow arrows highlight cells moving from the internal to the external compartment or vice versa. **J:** Classification of cell tracks according to their compartmental behavior during the 6-h imaging window. *“Exchange”* tracks undergo at least one excursion into the opposite compartment, whereas *“Stay”* tracks remain within their initial compartment. **K:** Duration, in minutes, of individual cell excursions into the opposite compartment. Scale bars: 100 μm (B), 50 μm (C, D, E), 25 μm (I). Dotted lines in C, D, E and I delineate the epithelium limit.

To examine whether this boundary was indeed static, we performed live imaging of embryonic explants and organoids expressing a nuclear marker (*NLS-mKate2*^18^) and a Notch activity reporter (*Hes1-emGFP*^19^) marking luminal commitment. While emGFP remained predominantly enriched in the inner compartment and excluded from the outer basal layer (Figure 1E; Figure S1H-J), the time-lapse movies revealed a surprisingly dynamic tissue; cells displayed extensive motility and frequently exchanged between the inner and outer compartments (Videos S5, S6). To characterize these behaviors, we next used automated nuclear segmentation and lineage tracking in organoids to reconstruct single-cell trajectories over 6-hour imaging windows (Figure S1K-M, Methods). Cells segregated into Hes1^high^ and Hes1^low^ populations (Figure 1F, G1, Methods); tracks displaying abrupt intensity fluctuations were confirmed to be tracking artifacts and excluded (Methods). The remaining Hes1^high^ and Hes1^low^ cell states remained stable throughout imaging (Figure 1H).

Single-cell trajectories quantified dynamic positional rearrangements within this globally segregated tissue organization. Individual Hes1^high^ and Hes1^low^ cells transiently crossed the compartment boundary, defined as a position within 5 µm from the basement membrane (BM), before returning to their original domain, both in organoids and in explants (Figure 1H, I; Video S7, 8). Over a 6-hour window, approximately 40% of Hes1^high^ cells and 20% of Hes1^low^ cells underwent at least one compartment exchange (Figure 1J; Methods). These excursions were short-lived, with a mean residence time of ∼40 minutes within the “incorrect” compartment before reversion (Figure 1K).

Together, these findings show that basal-luminal compartmentalization is maintained, during branching morphogenesis, as a dynamic, self-correcting steady state where misplaced cells are actively returned to their home domain, rather than as a static boundary.

### Differential adhesion and contractility maintain compartment boundaries through active cell repositioning

Having established that basal-luminal boundaries are maintained despite continuous inter-compartment cell exchanges, we next investigated the mechanism underlying this rapid positional restoration. Tissue boundaries can be stabilized when adjacent cell populations differ in their effective interfacial tension, an emergent property integrating cell-matrix adhesion, cell-cell adhesion, and actomyosin contractility^20,21^. We therefore asked whether lineage-specific adhesion-contractility states reinforce these compartments through a cell-sorting mechanism, consistent with a positional bias at their interface.

Across branched embryonic tissues, key mechano-modulators displayed striking compartment-specific localization. The laminin receptor Integrin *α*6 (ITGA6) was strongly concentrated at the outer, BM-facing interface (Figure 2A, B; Figure S2A, B). Actomyosin contractility markers, including Non-muscle Myosin heavy chain 2A (NMII-A, encoded by *Myh9*) and phosphorylated Myosin Light Chain (pMLC), were similarly enriched in the external basal compartment (Figure 2C, D; Figure S2C-F). In contrast, the adherens junction component E-cadherin (ECAD) was enriched within the internal luminal compartment (Figure 2E, F; Figure S2H, I), reflecting elevated intercellular cohesion. These spatial signatures were recapitulated in organoids (Figure 2G, H), indicating that basal and luminal cells retain distinct mechanical profiles outside the embryonic tissue context.

**Figure 2:**
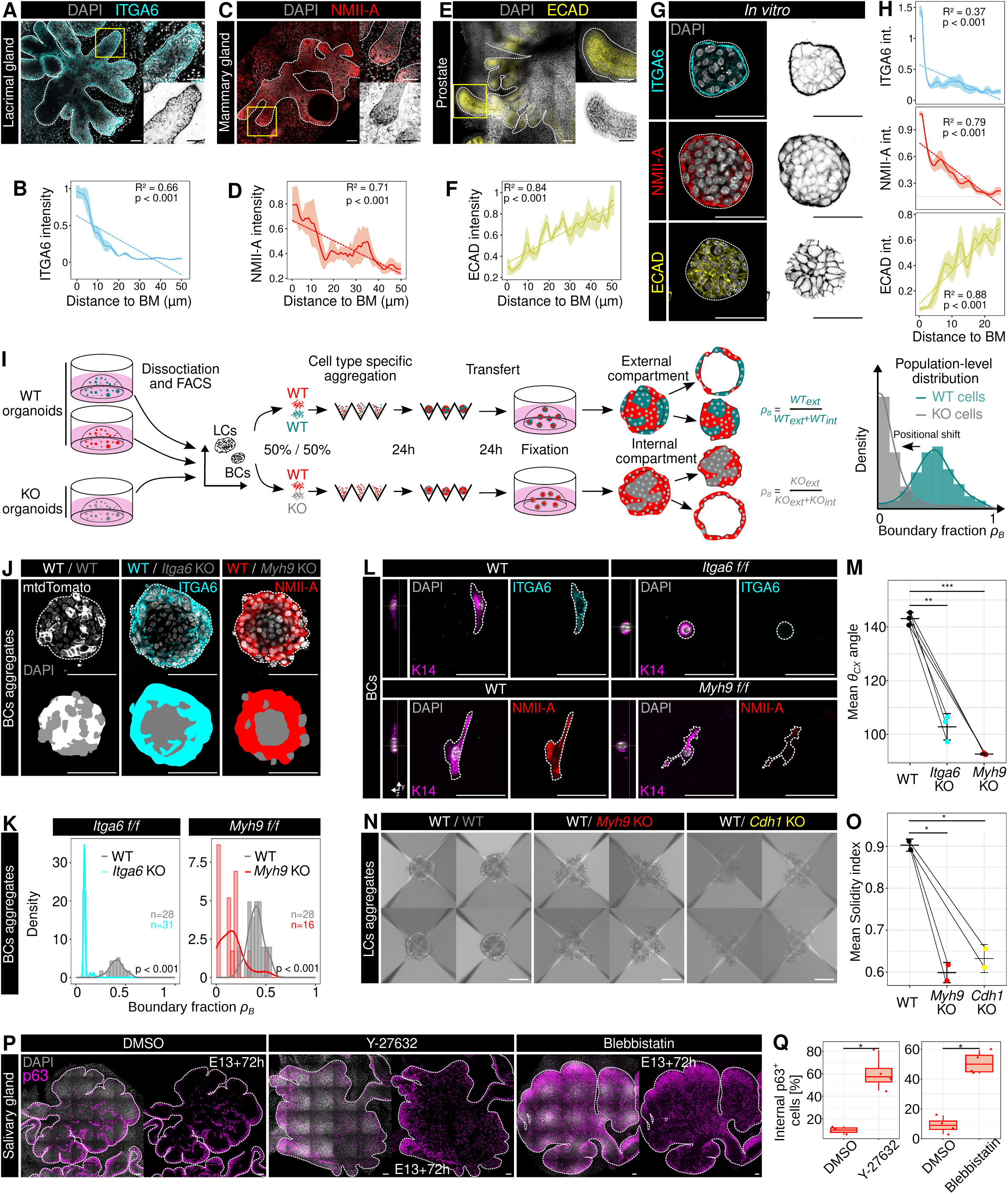
Adhesion-contractility organization regulates epithelial compartmentalization and cell positioning. **A:** Confocal imaging of ITGA6 in LG explants after 5 days in culture. **B:** Mean ITGA6 intensity profile as a function of distance from the BM in LG explants. **C:** Confocal imaging of NMII-A in MG explants after 7 days in culture. **D:** Mean NMII-A intensity profile as a function of distance from the BM in MG explants. **E:** Confocal imaging of ECAD in prostate explants after 5 days in culture. **F:** Mean ECAD intensity profile as a function of distance from the BM in prostate explants. **G:** Confocal imaging of ITGA6, NMII-A, and ECAD in mammary organoids after 7 days in culture. **H:** Mean ITGA6, NMII-A, and ECAD intensity profiles as a function of distance from the BM in organoids. **I:** Schematic representation of the workflow for generating epithelial aggregates from FACS-sorted mammary epithelial cells. Basal or luminal cells from control and knockout conditions were aggregated in AggreWell plates for 24 h and subsequently transferred to Matrigel for an additional 24 h. The quantification strategy is based on the spatial distribution of each population within the aggregate. Theoretical probability-density distributions illustrate the expected localization of WT (green) and KO (grey) cells. **J:** Confocal imaging of mixed basal aggregates containing WT and ITGA6- or MYH9-deficient cells after 4S h in culture. WT cells are identified by mtdTomato fluorescence. **K:** Probability-density distributions of WT (grey) and KO (cyan or red) cell positions within basal aggregates. Data were obtained from three biological replicates. **L:** Confocal imaging of WT, ITGA6-KO, and MYH9-KO basal cells plated on BME-coated substrates. **M:** Quantification of the mean cell–substrate contact angle (θ_cx_) for each genotype. Data were obtained from two biological replicates. **N:** Brightfield imaging of luminal aggregates composed of WT cells alone or mixtures of WT cells with MYH9-KO or ECAD-KO cells. **0:** Quantification of aggregate solidity, reflecting deviation from a compact spherical morphology, across the indicated conditions. Data were obtained from two biological replicates. **P:** Confocal imaging of p63 in salivary gland (SG) explants cultured for 3 days and treated during the final 2 days with Y-27632 (100 μM) or blebbistatin (20 μM). **Q:** Quantification of the percentage of p63-positive cells within the internal epithelial compartment following inhibition of actomyosin contractility. Scale bars, 50 μm. Dotted lines in A, C, E, G, J and P delineate the epithelium limit, and the cell membrane in N.

To test whether relative differences in adhesion or contractility between neighbouring cells are sufficient to bias their spatial positioning, we established same-lineage mosaic aggregates building on previous approaches using homotypic aggregates to isolate mechanical contributions to cell sorting^14,22^. This reductionist approach uncouples mechanical sorting from cell-identity signalling. Because mechanical sorting operates on a timescale of tens of minutes (Figure 1K), whereas basal lineage plasticity requires days (72-96 h^17^), assaying aggregates within 48 hours allowed us to isolate mechanical self-organization before fate conversions could occur.

We first generated basal mosaic aggregates containing wild-type (*WT/mtdTomato*) together with inducible knockout cells (*Itga6 f/f*^23^*, Cdh1 f/f*^24^ and *Myh9 f/f*^25^ driven by *R26-Cre^ERT^*^2 26^; Figure 2I). Loss of either ITGA6 or NMII-A consistently altered the boundary-fraction metric (*ρB*), consistently displacing mutant basal cells away from the BM-facing periphery and toward internal regions relative to their wild-type neighbours (Figure 2I, J, K; Figure S2J). Thus, altering cell-matrix adhesion or contractility within an otherwise common basal identity was sufficient to change the preferred spatial position of cells.

We next linked these sorting phenotypes to altered physical surface properties. In single-cell contact-angle assays on Matrigel-coated substrates, ITGA6- and NMIIA-deficient basal cells exhibited substantially reduced substrate affinity, displaying decreased contact angles (*θ*_CX_) comparable to wild-type luminal cells (Figure 2L, M; Figure S2K-M). Conversely, perturbations of NMII-A or ECAD in luminal cells strongly impaired intercellular cohesion, producing irregular and poorly compacted aggregates that precluded the radial sorting observed in basal aggregates (Figure 2N, O). Together, these experiments reveal complementary mechanical regimes in the two compartments: basal positioning depends strongly on cell-matrix adhesion and contractility, whereas luminal organization relies on cell-cell cohesion.

While these reductionist mosaic assays demonstrate that local mechanical disparities are sufficient to bias cell positioning, we asked whether active contractility is globally required to maintain compartment boundaries in intact, developing tissues. To test this, we acutely inhibited actomyosin contractility with blebbistatin or Y-27632 in *ex vivo* embryonic explants at developmental stages where basal and luminal compartments were already molecularly and spatially resolved. Global inhibition rapidly compromised boundary integrity across all four glandular tissues, triggering ectopic displacement of p63^+^ basal cells into internal domains (Figure 2P, Q; Figure S2N).

Together, these findings demonstrate that basal-luminal compartmentalization does not depend on static physical barriers. Instead, boundaries are actively preserved at steady state through lineage-specific adhesion and contractility, which provide a self-correcting mechanism that preserves tissue compartmentalization while permitting extensive cellular rearrangements.

### Emerging adhesion-contractility compartmentalization couples cell positioning to lineage allocation

Having established that relative mechanical disparities are sufficient to alter cell positioning in lineage-committed aggregates, we next asked whether introducing these same asymmetries prior to fate specification biases lineage output.

We began by examining the adhesion-contractility machinery at early developmental stages (MG/SG: E12, LG/Prostate: E15), where embryonic glandular epithelia consist predominantly of uncommitted multipotent progenitors. Across all four glandular tissues *in vivo* and in early organoid cultures, ITGA6, ECAD and NMIIA were initially homogeneously distributed (Figure 3A-E, S3A-H). As development progressed, these mechano-modulators became increasingly segregated, establishing spatially distinct mechanical domains (Figure 3A-E; S3A-H).

**Figure 3:**
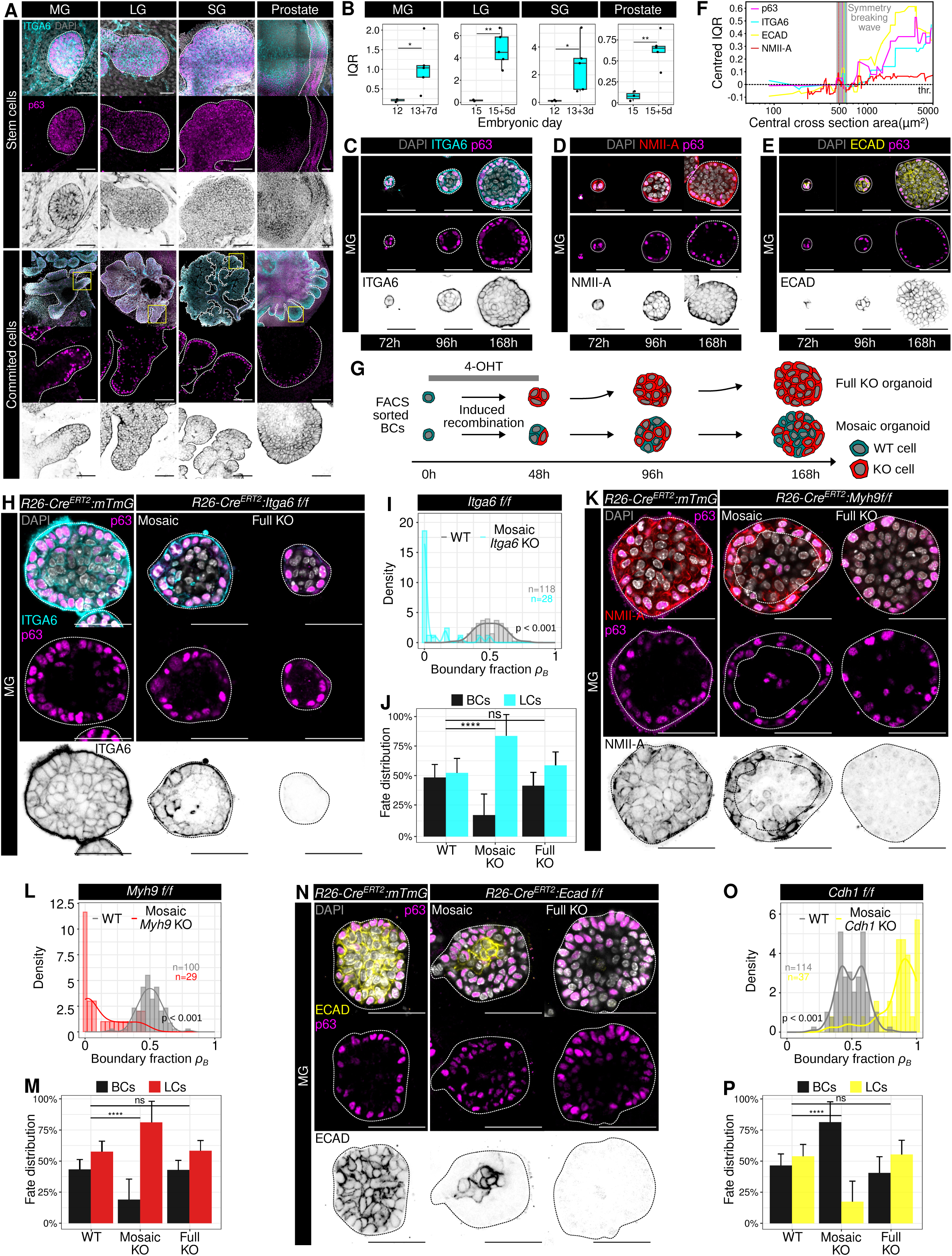
Emerging adhesion-contractility heterogeneity biases cell positioning and lineage fate through relative differences. **A:** Confocal imaging of ITGA6 and p63 in MG, SG, LG, and prostate embryonic explants at different developmental stages. Top: E12 (MG and SG), E15 (LG and Prostate). Bottom: E13+7d (MG), E13+3d (SG), E15+5d (LG and Prostate). **B:** Mean interquartile range (IQR) of cellular ITGA6 intensity across successive stages of embryonic development. Data were obtained from at least three biological replicates. **C-E:** Confocal imaging of p63 together with ITGA6, NMII-A, or ECAD, respectively, at successive stages of organoid growth. **F:** Centered IQR of p63, ITGA6, ECAD, and NMII-A expression as a function of organoid central cross section area, a proxy for organoid cell number, relative to the window of fate symmetry breaking. **G:** Schematic representation of the workflow used to generate fully recombined or mosaic knockout organoids from FACS-sorted mammary BCs. Single BC-derived organoids were exposed to OHT during the first 4S h of culture. Complete recombination generates fully mutant organoids, whereas partial recombination generates mosaic organoids containing both WT and mutant cells. **H:** Confocal imaging of ITGA6 in WT, mosaic, and fully recombined ITGA6-KO organoids. **I:** Measured probability-density distribution of ITGA6-KO cell positions within mosaic organoids. **J:** Fate distribution of WT and ITGA6-KO cells in mosaic and fully mutant contexts. **K:** Confocal imaging of NMII-A in WT, mosaic, and fully recombined MYH9-KO organoids. **L:** Measured probability-density distribution of MYH9-KO cell positions within mosaic organoids. **M**: Fate distribution of WT and MYH9-KO cells in mosaic and fully mutant contexts. **N:** Confocal imaging of ECAD in WT, mosaic, and fully recombined ECAD-KO organoids. **0:** Measured probability-density distribution of ECAD-KO cell positions within mosaic organoids. **P:** Fate distribution of WT and ECAD-KO cells in mosaic and fully mutant contexts. Scale bars, 50 μm. Dotted lines in A, C, D, E, H, K, and N delineate the epithelium limit.

To quantify the onset of lineage segregation, we measured the interquartile range (IQR) of immunostaining intensities across individual cells over developmental time (Figure 3B, F; Figure S3D, E, I-L; Methods)^17^. Because the IQR captures population-wide expression variance as interior cells diverge from surface cells, this metric reliably identifies symmetry breaking. The sharp increase in adhesion-contractility heterogeneity coincided with the emergence of bimodal p63 expression at the ∼13-cell stage (500 µm^2^), a transition previously shown to mark initial lineage specification^17^ (Figure 3F; S3I-L). Mechanical compartmentalization therefore arises concomitantly with fate commitment across branching epithelia.

Because compartment-specific mechanical properties are sufficient to bias cell positioning (Figure 2), and spatial position dictates exposure to niche-derived fate cues^17^, we reasoned that introducing local mechanical disparities during early development would bias lineage commitment by altering niche occupancy. To test this, we induced Cre-mediated recombination in single basal cell-derived organoids to generate either uniform knockouts or mosaic organoids containing an intermixture of wild-type and mutant cells (*Itga6 f/f*, *Myh9 f/f*, or *Cdh1 f/f*; Figure 3G). Strikingly, uniform deletion of these regulators did not impair lineage specification. Organoids maintained basal and luminal proportions comparable to wild-type controls, demonstrating that these adhesion-contractility components are dispensable for cell-autonomous differentiation (Figure 3H, J, M, P).

In mosaic organoids, however, mutant cells exhibited pronounced, coordinate shifts in both positioning and fate relative to their wild-type neighbours (Figure 3H-P; Figure S3M, N). Loss of ITGA6 or NMII-A displaced mutant cells into the internal compartment and significantly increased their contribution to the luminal lineage. Conversely, ECAD depletion biased cells toward the external compartment and enriched basal identity (Figure 3H-P). Thus, rather than directly dictating fate programs, relative mechanical disparities determine the probability with which cells access lineage-instructive spatial niches.

These findings demonstrate that adhesion-contractility states bias lineage commitment non-autonomously through positional allocation, raising the question of when and where fate transitions occur as cells relocate between compartments.

### Pre-commitment niche residency predicts lineage specification

The contrasting phenotypes of mosaic versus uniform knockouts indicate that relative mechanical disparities bias lineage decisions by directing cells into distinct spatial niches. We therefore asked how cell movements across boundaries prior to fate specification were linked to definitive lineage commitment and whether a cell positional history before commitment could predict its future lineage. To address this, we reconstructed single cell lineage trees from early organoid growth through symmetry breaking using long-term live imaging of *NLS-mKate2:Hes1-emGFP* organoids (Figure 4A; Video S9, Methods). Organoids followed a stereotyped developmental trajectory: successive near-synchronous division waves expanded the epithelium from 8 to 16 and 32 cells (Figure 4B). Lineage symmetry breaking, defined as the emergence of *Hes1-emGFP* heterogeneity, consistently occurred during the division wave transitioning from 8 to 16 cells (Figure 4C, D; Methods). This predictable schedule allowed us to register independent organoid movies to this 8-to-16-cell transition for retrospective tracking (Figure S4A).

**Figure 4:**
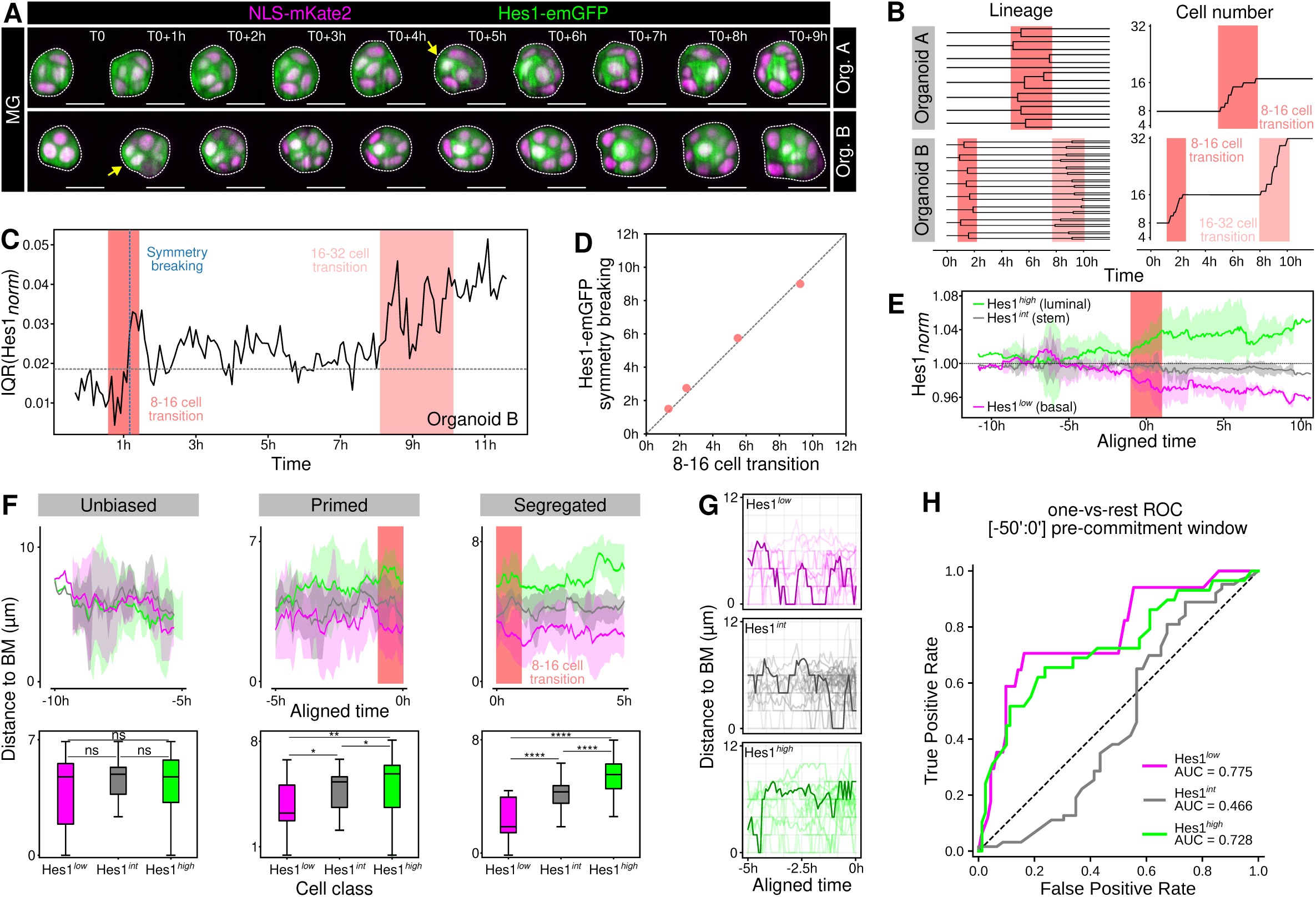
Pre-commitment cell positioning predicts HE51 symmetry breaking and lineage stabilization. **A:** Representative time-lapse optical sections of organoids expressing the nuclear reporter NLS-mKate2 and the HES1 reporter Hes1-emGFP. Yellow arrows indicate symmetry-breaking events, marked by the emergence of cells with increased Hes1-emGFP levels relative to the initial homogeneous state. Images were extracted from live imaging covering a window between 4Sh and 120h post seeding. **B:** Lineage trees and corresponding cell-number trajectories for the two organoids shown in A. Shaded regions indicate the S-16- and 16-32-cell transitions. **C:** Interquartile range of normalized Hes1-emGFP intensity across cells over time in organoid B. The onset of HES1 symmetry breaking and the S-16- and 16-32-cell transitions are indicated. **D:** Relationship between the timing of the S-16-cell transition and the onset of HES1 symmetry breaking across individual organoids. The dashed line indicates equality between the two timings. **E:** Mean normalized Hes1-emGFP intensity over time for cells classified as Hes1^high^, Hes1^int^, or Hes1^low^, aligned to the S-16 transition. Shaded areas indicate variability between cells. **F:** Distance of Hes1^low^, Hes1^int^, and Hes1^high^ cells from the BM during successive temporal windows preceding and following HES1 symmetry breaking, corresponding to unbiased, primed, and biased positional states. Boxplots below summarize cell-BM distances for each HES1 class within each window. **G:** Individual cell trajectories showing distance from the BM during the pre-commitment window, stratified by subsequent Hes1^low^, Hes1^int^, and Hes1^high^ classification. **H:** One-versus-rest ROC analysis testing the ability of cell position during the pre-commitment window (-5 to 0 h) to distinguish subsequent HES1 classes. AUC values are indicated for each class. Scale bars, 50 μm. Dotted lines delineate the epithelium limit.

In contrast to the mature stages analyzed in Figure 1, where *Hes1-emGFP* predominantly demarcate cells into stable Hes1^high^ (luminal) and Hes1^low^ (basal) populations, early organoids exhibited overlapping signals. Prior to symmetry breaking, cells occupied a continuum of states comprising intermediate *Hes1-emGFP* levels: Hes1^high^ (luminal-like), Hes1^low^ (basal-like) or Hes1^int^ (stem cell-like/intermediate) (Figure 4E; Methods).

We next examined the positional history of individual cells leading up to fate segregation, defining three distinct temporal windows: an unbiased phase (>5 h before symmetry breaking), an emergent primed phase (the final ∼5 h prior to symmetry breaking), and a segregated phase (post-symmetry breaking) (Figure 4F). At the population level, distance to the basement membrane (BM) did not differ among future fate classes during the unbiased window 5 to 10 hours prior to symmetry breaking (Figure 4F). However, during the emergent bias window (the final ∼5 h), cells destined for a Hes1^high^ luminal fate localized significantly farther from the BM than prospective Hes1^low^ and Hes1^int^ cells, an asymmetry that was amplified upon symmetry breaking (Figure 4F). Despite this diverging population average, individual cell trajectories exhibited extensive wander and positional overlap (Figure 4G), indicating that instantaneous position at any single timepoint cannot reliably predict fate at single-cell resolution.

We therefore asked whether integrating a cell’s positional history over time would be more informative by capturing cumulative niche exposure. Spearman correlation between time-integrated distance to BM and final fate peaked within a narrow window approximately 50 minutes prior to symmetry breaking (ρ = 0.427, permutation p = 5×10^−5^) whereas earlier windows showed a substantially weaker association (Figure S4B, C). A one-versus-rest ROC analysis confirmed that pre-commitment residency was predictive for both committed lineages; cells destined for a luminal fate (Hes1^high^) spent significantly more time away from the basement membrane than their peers (AUC = 0.728), while future basal cells (Hes1^low^) spent more time proximal to the BM (AUC = 0.775; Figure 4H). In contrast, Hes1^int^ were not distinguished by residency (AUC = 0.466), reflecting continued niche sampling without stable positional residence.

Together, these findings demonstrate that lineage commitment does not depend on instantaneous position but is predicted by cumulative niche residency within a critical pre-commitment window.

### Lineage-regulatory pathways tune adhesion-contractility states and epithelial dynamics

Having shown that relative mechanical perturbations bias cell positioning and lineage output, we next examined the opposite scenario: whether lineage-regulatory pathways reciprocally modulate the adhesion and contractility states underlying tissue compartmentalization. We first perturbed lineage regulatory pathways from the onset of organoid formation and quantified by immunofluorescence the intensity and distribution of ITGA6, ECAD, and NMIIA. Pharmacological Notch inhibition (DAPT), which prevents luminal commitment^17^, increased ITGA6 and NMIIA global intensity without substantially altering the relative differences between external and internal compartments (Figure 5A-C). Orthogonal validation via flow cytometry and *ex vivo* explants confirmed a global increase in ITGA6 levels following DAPT treatment (Figure S5A-C). Conversely, Notch activation during early stage of organoid growth, known to promote luminal fate commitment^17^, reduced ITGA6 and NMIIA levels while increasing ECAD intensity in mutant cells (Figure 5D-F). Extending beyond Notch signalling, both pharmacological and genetic activation of YAP, which maintains stem-like progenitor identity^17^, selectively reduced NMIIA levels in the external compartment (Figure 5G-I), without affecting E-cadherin or ITGA6 levels (Figure S5D-F).

**Figure 5:**
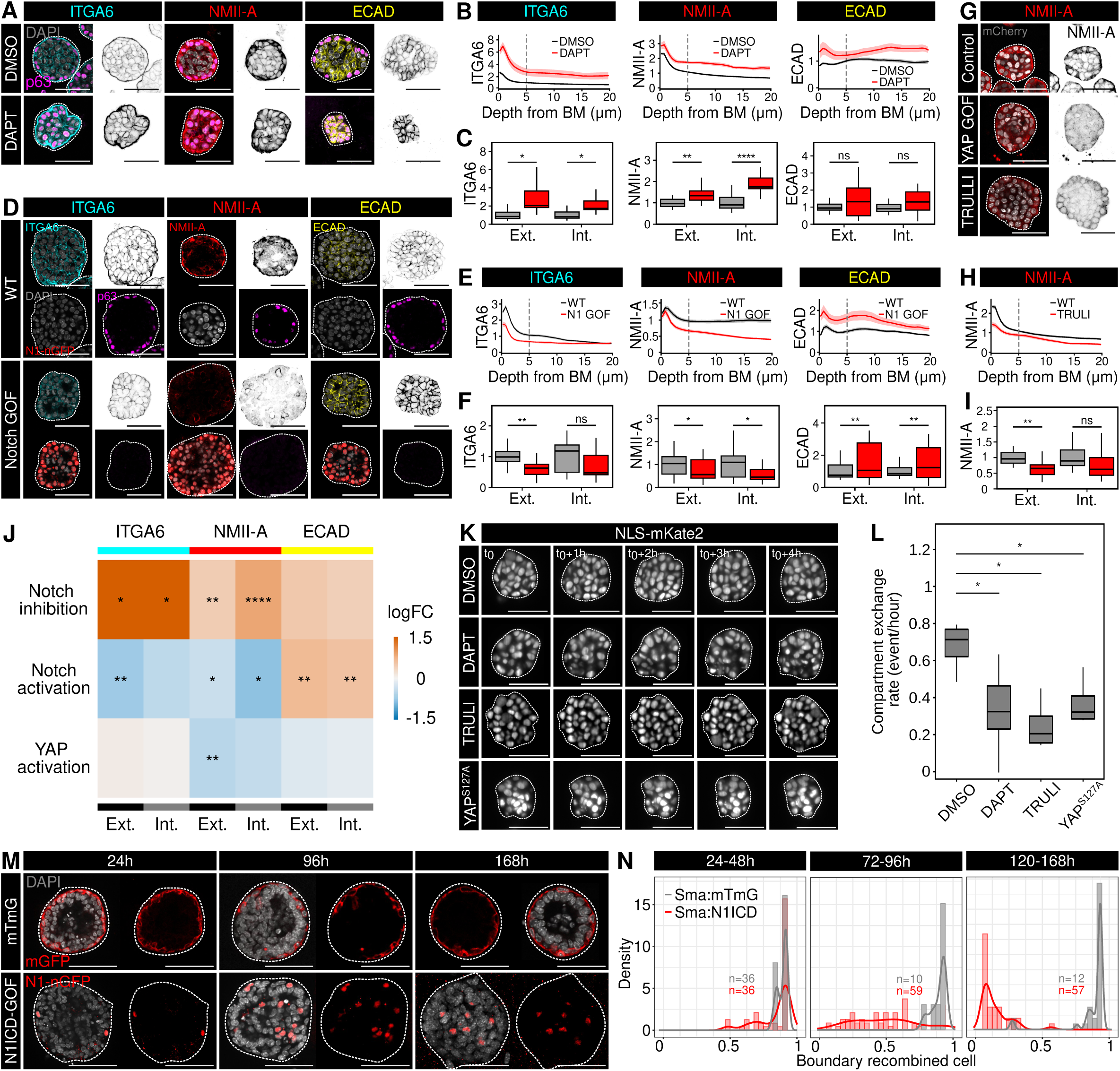
Lineage-regulatory pathways remodel adhesion-contractility organization and epithelial dynamics. **A:** Confocal imaging of ITGA6, NMII-A, and ECAD in organoids treated with DMSO or with 10 μM of the Notch inhibitor DAPT. **B:** Mean intensity profiles of ITGA6, NMII-A, and ECAD as a function of distance from the basement membrane (BM) in DMSO- and DAPT-treated organoids. **C:** Quantification of ITGA6, NMII-A, and ECAD levels in the external (Ext.) and internal (Int.) compartments following Notch inhibition. **D:** Confocal imaging of ITGA6, NMII-A, and ECAD in control (WT) and Notch gain-of-function (Notch1-GOF) organoids. **E:** Mean intensity profiles of ITGA6, NMII-A, and ECAD as a function of distance from the BM in WT and Notch1 GOF organoids. **F:** Quantification of ITGA6, NMII-A, and ECAD levels in the external and internal compartments of WT and N1 GOF organoids. **G:** Confocal imaging of NMII-A in control organoids and following YAP activation, either genetically (YAP GOF) or pharmacologically with TRULI. **H:** Mean NMII-A intensity profile as a function of distance from the BM in control and TRULI-treated organoids. **I:** Quantification of NMII-A levels in the external and internal compartments following YAP activation. **J:** Heatmap summarizing changes in ITGA6, NMII-A, and ECAD levels in the external and internal compartments following Notch inhibition, Notch activation, or YAP activation. Values are shown as log fold changes relative to the corresponding control condition. **K:** Representative time-lapse optical sections of mammary organoids expressing the nuclear reporter *NLS-mKate2* under control conditions, following Notch inhibition with DAPT, pharmacological YAP activation with TRULI, or genetic YAP activation by *YAP^S^*^127^*^A^*. **L:** Quantification of compartment-exchange frequency, expressed as the number of exchange events per hour, under the indicated conditions. **M:** Confocal imaging of control *Sma-Cre^ERT^*^2^*:mTmG* and *Sma-Cre^ERT^*^2^*:N1/CD-GOF* mammary organoids at 24, 96, and 16S h of culture, showing the spatial distribution of recombined cells over time. **N:** Probability-density distributions of the position of recombined cells relative to the organoid boundary during the indicated time windows (24-4S h, 72-96 h, and 120-16S h) in control and N1ICD-GOF organoids. Scale bars, 50 μm. Dotted lines in A, D, G, K and M delineate the epithelium limit.

Together, these findings indicate that lineage-regulatory pathways modulate specific adhesion and contractility components (Figure 5J).

We next asked whether these changes in cell identity and adhesion and contractility states were accompanied by alterations in epithelial rearrangement dynamics. Live imaging revealed that DAPT treatment reduced cell exchanges between spatially defined external and internal compartments (Figure 5K, L; Video S10). Constitutive YAP activation had an even more pronounced effect, largely immobilizing organoid cells and suppressing inter-compartment transitions despite continued proliferation (Figure 5L; Videos S11, S12). Lineage homogenization therefore arrests dynamic neighbour exchanges, suggesting that active tissue remodeling relies on fate heterogeneity across the epithelium.

Finally, we investigated whether changing the identity of individual cells within a mosaic epithelium could redirect their positioning toward the compartment corresponding to their newly acquired fate. To test this, we used mosaic Notch activation to induce luminal differentiation, as previously established *in vivo*^27^ and in organoids^28^. As expected, when recombination was induced in stem cells, N1ICD-expressing cells preferentially occupied the organoid interior (Figure S5G, H). Notably, Notch activation in already committed basal cells at the organoid surface promoted luminal conversion accompanied by inward repositioning (Figure 5M, N). Lineage-regulatory signalling pathways therefore reshape adhesion-contractility profiles, regulate tissue dynamics, and autonomously drive cell sorting toward fate-appropriate compartments.

## Discussion

Developing glandular epithelia preserve sharply segregated basal and luminal compartments despite rapid growth, branching morphogenesis and extensive cell rearrangements. Through long-term live imaging, we have demonstrated that this compartmentalization does not depend on rigid physical confinement. Instead, basal and luminal cells frequently cross the compartment interface, yet long-term mispositioning remains rare. Previous studies have established that differences in cell adhesion and actomyosin contractility contribute to cell segregation and the formation of developmental boundaries^5,10–12,28^. However, conventional endpoint analyses cannot distinguish whether these mechanical differences primarily restrict cell crossing or favor the restoration of appropriate positioning following transient exchanges. In branching glandular epithelia, our findings demonstrate that lineage-specific mechanical profiles tolerate continuous cell exchanges while actively enforcing sorting. We therefore propose the existence of a permissive, self-correcting boundary where local interfacial crossings are permitted, but relative mechanical differences between neighbouring cells continuously bias them toward their native compartment. Whether cells acquire specialized mechanical properties at heterotypic basal-luminal contacts, as described in other developmental boundaries^11,12^, remains an important question for future work.

Although mitotic rounding has been proposed to challenge tissue compartmentalization^7^, our observations indicate that cell division alone cannot account for the frequent basal-luminal exchanges observed in embryonic tissues and organoids. Indeed, constitutive YAP activation preserved high proliferation while virtually abolishing inter-compartment transitions, demonstrating that epithelial rearrangement dynamics are regulated independently of cell cycle progression. Such cellular mobility may be indispensable during branching morphogenesis, when tissues must undergo extensive remodeling while preserving their compartmental organization. Consistent with this view, differences in cell-matrix and cell-cell adhesion between peripheral and internal cells facilitate epithelial remodeling and folding during salivary gland branching morphogenesis^16^. Glandular epithelia may therefore achieve robustness through error correction rather than rigid error prevention, preserving morphogenetic flexibility while maintaining tissue-scale compartmentalization. This strategy mirrors physical models wherein developmental robustness emerges from tolerating local fluctuations that are subsequently resolved by self-organizing feedback mechanisms^30,31^. Whether this mode of boundary maintenance is specifically deployed during periods of extensive tissue remodeling, such as development, puberty or pregnancy or simply operates on extended timescales in mature tissues remains an open question.

Beyond maintaining compartmental organization, these positional dynamics may influence lineage commitment by dictating the microenvironments that cells experience. Mechanical properties are known to influence cell fate both through direct mechano-transduction and by regulating cell sorting and positioning, notably during early embryonic development^32^. Our mosaic perturbations suggest that a similar coupling operates in glandular epithelia, with a critical role for the mechanical properties of neighbouring cells. Altering adhesion or contractility components biases both cell positioning and lineage allocation when mutant cells coexist wild-type neighbours, yet these fate biases are completely lost when identical perturbations are applied homogeneously across the entire tissue. This context dependence suggests that relative mechanical states, rather than a given mechanical signalling alone, dictates lineage choices by directing spatial niche residency. Conversely, manipulating Notch or YAP remodels compartment-specific adhesion and contractility and alters cell-rearrangement dynamics, indicating that lineage decisions reciprocally establish the physical properties required for sorting. Such reciprocal coupling is not unique to glandular tissue. In intestinal organoids, WNT, Notch and Hippo-dependent lineage patterning generates spatially distinct stem cell and differentiated enterocyte domains with regional differences in actomyosin contractility that contribute to crypt formation^33^. Mechanical compartmentalization thus represents a general organizing principle of branching and folded epithelia, whereby emerging transcriptional states generate local physical disparities that coordinate spatial sorting.

While temporal niche residency is a recognized hallmark of stem cell maintenance, its role in developing glandular epithelia fundamentally redefines classical paradigms. In classical niches, such as the intestinal crypt^34^ and the *Drosophila* ovarian germline^35^, residence within a spatially restricted microenvironment preserves stemness, whereas displacement triggers differentiation. Developing glandular epithelia, however, lack an equivalent localized niche. Instead, our findings indicate that the emerging basal and luminal compartments function as mobile, lineage-stabilizing niches in which prolonged residency consolidates fate commitment via sustained exposure to compartment-specific biochemical and mechanical cues. Across both systems, niche residency thus operates as a universal state-stabilizer linking positional history to fate, though the nature of that state depends on the local signal. While the fluorescent protein stability of our HES1 reporter does not allow us to identify rapid transcriptional resets, our model predicts that cells undergoing repeated compartment switching will retain developmental plasticity longer than persistently resident cells. Dissecting the direct causal interplay between positional history and the kinetics of fate restriction will be an exciting direction for future studies.

Synthesizing these findings leads us to propose that branching tissues safeguard compartmental fidelity through two coupled, timescale-separated error-correcting mechanisms. On a rapid kinetic timescale (tens of minutes), interfacial tension and differential adhesion act as a mechanical filter, driving transiently displaced cells back to their native compartment before their identity is altered. On a slower timescale (hours to days), cells that escape this mechanical correction and remain stranded in the opposing niche could undergo lineage reprogramming, adopting the fate of their new microenvironment to purge rogue identities from the tissue.

Finally, the conservation of these dynamic boundary mechanisms across glandular epithelia of divergent embryological origins suggests that this strategy represents a shared evolutionary solution to the physical challenges of branching morphogenesis. We previously identified conserved signalling principles underlying fate specification across these tissues^36^; the present findings suggest that this conservation may extend to the mechanical rules that couple cell identity to spatial organization. More broadly, this raises the possibility that evolution conserves not only molecular signalling pathways, but higher-order physical and organizational principles that make tissue architecture robust to the cellular rearrangements inherent to morphogenesis.

## Resource availability

### Material availability

This study did not generate new unique reagents.

### Data and code availability

All data supporting the conclusions can be found in the main text or are available upon request from the corresponding authors, Robin Journot and Silvia Fre. The raw data and the code used for image analysis will be available on zenodo upon publication. Any additional information required to analyze the data reported in this paper is available from the corresponding authors upon request.

## Acknowledgments

We wish to acknowledge all Fre team members, as well as our colleagues Dr. Yohanns Bellaiche, Dr. Edouard Hannezo and Prof. Michel Labouesse, for technical advice and constructive discussions. The authors acknowledge the Cell and Tissue Imaging Platform PICT-IBiSA, member of the national infrastructure France-Bioimaging (https://ror.org/01y7vt929), supported by the French National Research Agency (ANR-24-INBS-0005 FBI BIOGEN) and the Cell and Tissue Imaging (PICT-IBiSA), Institut Curie, member of France-Bioimaging of the Genetics and Developmental Biology Unit (UMR3215/U934), as well as the In Vivo Experimental Facility, particularly Cedrick Pauchard, Mickael Garcia and Celine Daviaud, for help in the maintenance and care of our mouse colony. Authors wish to acknowledge our colleagues Dr. David Rozema and Dr. Peta Bradbury for their help with LS1 microscope and Dr. Anna-Katerina Hadjantonakis (Developmental Biology Program - Sloan Kettering Institute Memorial Sloan Kettering Cancer Center) for agreeing to share the NLS-mKate2 mice with us. We want to thank Dr. Clementine Villeneuve for their critical feedback during the preparation of the manuscript.

This work was funded by, the French National Research Agency (ANR) grant numbers ANR-21-CE13-0047, ANR-22-CE13-0009 and ANR-24-CE13-3529-01, the Medical Research Foundation FRM “FRM Equipes” EQU201903007821, the FSER (Fondation Schlumberger pour l’éducation et la recherche) FSER20200211117, the Association for Research against Cancer (ARC) label ARCPGA2021120004232_4874, the Worldwide Cancer Research Foundation # 24-0216, the Ligue contre le Cancer label ELF_FRE_26738 and by Labex DEEP ANR-Number 11-LBX-0044.

The Novo Nordisk Foundation Center for Stem Cell Medicine is supported by the Novo Nordisk Foundation (NNF21CC0073729).

R.J. was funded by a PhD fellowship from the French minister of research and by the FRM (FDT202404018370). R.J. received an EMBO Scientific Exchange Grant (11124) to perform part of the experimental work.

The funding agencies had no role in study design, data collection and analysis, decision to publish, or preparation of the manuscript.

## Author Contributions

R.J. and S.F. conceived and designed the experiments. R.J., T.D., L.H., V.S., C.M., A.B. and C.C. performed all experiments. M.M.F. contributed to experimental work and edited the manuscript. P.A.S., M.M., and K.B.J., provided *R26-rtTA:tetO-YAP^S^*^127^*^A^* and *R26-rtTA:tetO-mCherry* samples and hosted part of the experimental work. J.L.M. provided the *Cdh1* f/f and *Myh9* f/f mouse lines and access to the LS1 microscope. R.J. curated, interpreted results and prepared the figures. R.J, and S.F. wrote the manuscript. V.S. and Z.J.G. provided important conceptual input during manuscript finalization. S.F. provided funding, project administration and supervision. All authors reviewed and approved the manuscript.

## Declaration of interests

The authors declare no competing interests.

## Methods

### Ethics statement

All studies and procedures involving animals were carried out in strict accordance with the recommendations of the European Community (2010/63/UE) for the protection of vertebrate species used for experimental and other scientific purposes. Approval was granted by the Ethics Committee of the Institut Curie CEEA-IC #2021-029 and the French Ministry of Research (reference #34364-202112151422480). We adhere to the internationally established principles of Replacement, Reduction and Refinement (3Rs) according to the Guide for the Care and Use of Laboratory Animals (NRC 2011). Animal husbandry, supply, maintenance and care in the animal facility of the Institut Curie (facility license #C75-05-18) before and during experiments fully met the needs and welfare of the animals. All animals were housed in individually ventilated cages under a 12:12h light/dark cycle, with water and food available *ad libitum*. All mice were culled by cervical dislocation. Mice were genotyped by PCR on genomic DNA extracted from an earpiece for adult mice or tail tip for embryos. Mouse breeding and husbandry was managed using the mouse colony organization software MiceManager: https://infenx.com/mouse-colony-management-software/.

### Animal models

Organoids were derived from 8-12 weeks male and female from the following lines: *R26-CreERT2*^26^ or *Sma-CreERT2*^37^ mouse crossed with *R26:mTmG*^38^, *Cdh1 f/f*^24^*, Myh9 f/f*^25^*, Itga6 f/f*^23^ or *N1CD-nGFP*^19^ animals. *Rosa26-M2rtTA*^39^ (Jackson stock # 006965) crossed with *tetO-YAP S127A*^40^ and *tetO-H2B:mCherry*^41^ (Jackson stock # 014602). All mice used were in a mixed genetic background. Detailed description *of in vitro* organoid cultures is provided in the Method details section.

We crossed the *R26-Cre^ERT^*^2^ line with *mTmG*, *Hes1-emGFP*^19^ or *R26:NLS-MKate2*^18^ (Accession N°: CDB1283K) for embryonic experiments. All mice used were in a mixed genetic background. Reporters and mutant alleles were recombined by a single intraperitoneal injection of tamoxifen free base prepared in sunflower oil containing 10% ethanol (0.1 mg per g of mouse body weight), unless indicated otherwise. The explants were collected on embryo at embryonic day (E) 13 for mammary and salivary gland, and at E15 for lacrimal gland and prostate. Detailed description of *ex vivo* organoid cultures is provided in the Method details section.

### Replicate definition

Experiments were performed in biological and technical replicates as indicated. A biological replicate corresponds to pups from different litters for embryonic experiments. Siblings included in the same experiment are considered technical replicates.

### Statistics, reproducibility and data availability

Reproducibility codes for the analyses are available online via GitHub, as detailed in the ‘Code availability’ section. All statistical methods are described in the corresponding sections of the paper. Atlases of individual tissues and integrated dataset will be shared with associated metadata upon publication. Experiments were performed in biological replicates as indicated. A biological replicate corresponds to different animals. Experiments with at least 3 biological replicates were used to calculate the statistical value of each analysis. All graphs are mean + SD. Statistical analyses were performed using the Wilcoxon test or Chi squared test and are stated in the corresponding figure legends. *: p-value < 0.05, **: p-value < 0.01, ***: p-value <0.001, ****: p-value < 0.0001.

### Organoid line culture establishment

MG, LG, SG and prostate were isolated from 8-12-week-old mice. A typical experiment was generated using two males and two females, with MG and Prostate from two animals pooled, while LG and SG from four animals were pooled and the four tissues were dissociated using the same protocol as previously described^42^. Briefly, single cell dissociation was performed by enzymatic digestion with 600 U/ml of collagenase for 90 min at 37°C with shaking (160 rpm). Cells were further dissociated in hot trypsin for 1 min, in 5 mg/mL dispase and 0.1 mg/mL DNase I for 3 min at 37°C, then red blood cells were eliminated in 0.63% NH_4_Cl. To derive organoid lines, resulting fragments were directly embedded in Matrigel at a density of 1000 fragments per 10 μL BME. The primary organoids were cultured in media detailed below.

### Flow cytometry and sorting

To isolate single cells from organoid lines, organoids were extracted from Matrigel dome using cold PBS and then Cell Recovery solution on ice for 10 minutes. Cells were then dissociated in hot trypsin for 1 min, in 5 mg/mL dispase and 0.1 mg/mL DNase I for 3 min at 37°C. The cell suspension was filtered through a 40μm cell strainer to obtain a single cell preparation for FACS. Single cell suspensions were incubated for 30 minutes on ice with conjugated primary antibodies diluted in PBS (APC anti-mouse CD31, APC anti-mouse Ter119, APC anti-mouse CD45, APC/Cy7 anti-mouse CD49f (primary cells) or Pdpn (organoid line)^43^, PE/Cy7 anti-mouse EpCAM at 1:100 dilution). Cells were washed with PBS, resuspended in flow buffer containing DAPI and passed through a 40 μm cell filter. FACS analysis was performed on ARIA flow cytometer (BD). Doublets, dead cells (DAPI+) and Lin+ (CD45+, CD31+, Ter119+) cells were systematically excluded from the analysis. Full details of the antibodies used can be found in Table S1. Results were analyzed using FlowJo software (V10.0.7). These cells were then expanded as organoid lines.

To isolate and sort organoid cells, two wells of organoids were harvested in ice-cold PBS and the pellets were then incubated on ice for 15 minutes in cold Cell Recovery Solution to dissolve the Matrigel. Organoids were then centrifuged at 400 × g for 5 minutes at 4°C. Cell dissociation was performed as previously described for tissues.

### *In vitro* organoid cultures

Sorted BCs were embedded in 90% Matrigel and cultured in 24-well plates or 8-well imaging chambers (Ibidi 80827) for whole mount staining. After polymerization, the Matrigel drops were covered with tissue-specific organoid medium containing a base of DMEM/F12 supplemented with 1x Glutamax, 1x HEPES, 1X B27 supplement, recombinant Noggin and R-spondin, stabilized FGF2 and 2% penicillin/streptomycin. MG medium was supplemented with 1x N2 supplement and Neuregulin, SG and LG medium were supplemented with NAC, Wnt3a and EGF and prostate medium was supplemented with dihydrotestosterone (DHT), EGF and A8301 ^44–47^. All organoid cultures were supplemented with Y27632 for the first 48 hours of culture. Factors and inhibitors are listed in Table S2.

### Aggregate formation

Basal and luminal cells were isolated by FACS as described above and seeded into 24-well AggreWell 400 (34415) plates according to the manufacturer’s instructions. Aggregates were generated with a target size of 100 cells, by combining 50 cells from each population. For mosaic experiments, the two populations corresponded to WT and mT/mG-reporter-labelled cells, or WT and mutant cells, depending on the experiment. Cells were allowed to aggregate before further culture and analysis as indicated for each experiment.

### Contact-angle measurements

FACS-isolated basal cells (BCs) and luminal cells (LCs) were seeded onto glass-bottom Ibidi wells coated for 1 h with 1% basement membrane extract (BME). Cells were allowed to attach overnight, washed to remove non-adherent cells, fixed, and processed for immunofluorescence staining. Samples were subsequently imaged by confocal microscopy for contact-angle analysis.

### Generation and isolation of knockout cells

To obtain organoid cultures with a high proportion of recombined cells, primary-derived mutant organoid lines were expanded for at least two passages in medium supplemented with 500 nM 4-hydroxytamoxifen (4-OHT) to induce Cre-mediated recombination. Organoids were subsequently dissociated and mutant cell populations isolated by FACS using the procedures described above.

### Embryonic tissue isolation and explant culture

Embryonic tissues were collected at two different stages of development. Mammary and salivary buds were collected at E13, while the urogenital sinus (UGS) and lacrimal gland were collected from E15 embryos as previously described^17^. Briefly, embryos were collected from the uterus of a pregnant female and stored on ice in 24-well plates. Embryos were genotyped for sex and potential transgene. Dissection of SG, LG and UGS proceeded directly, but embryos were kept in PBS at 4°C overnight before dissection of MG buds. The epidermis covering the flank and MG of the embryos was removed without enzymatic digestion.

Embryonic tissues were then placed on a cell culture insert, floating on culture medium, in 6-well plates. The embryonic culture medium contains a base of DMEM/F-12 supplemented with 1x Glutamax, 1x HEPES and 2% penicillin-streptomycin. The MG medium is supplemented with 10% fetal bovine serum (FBS) (v/v). The LG and SG medium are supplemented with 2% insulin-transferrin-selenium (ITS), while the UGS (prostate) medium also contains 10 µM dihydrotestosterone (DHT) (See Table S2). Explant cultures grew at 37°C with 5% CO2 and the medium was replaced every 48 hours.

### Image acquisition

Images were acquired using an LSM780, LSM880 or LSM900 inverted laser scanning confocal microscope (Carl Zeiss) equipped with 25x/0.8 OIL LD LCI PL APO or 40x/1.3 OIL DICII PL APO objectives. For embryonic explants, images were acquired using a LSM780 or LSM880 inverted laser scanning confocal microscope (Carl Zeiss) equipped with 40x/1,0 W DIC PL APO VIS-IR (441452-9900) objectives. Z-stack were acquired every 5 minutes with a X, Y resolution of (0.512×0.512 µm) and a Z resolution of 2 µm. The organoid time-lapse were acquired on a Viventis LS1 light-sheet microscope Z-stack were acquired every 5 minutes with a X, Y resolution of (0.512×0.512 µm) and a Z resolution of 1 µm. All quantitative imaging analyses were performed in Python.

### Immunostaining

Whole mount immunostaining was performed on organoids and explants. Organoids were fixed in ice-cold 4% PFA (10 minutes), permeabilized with PBS 1% Triton x100 (1 hour) for organoids and with PBS 5% Triton x100 (24 hours) for explants, blocked in PBS, 5% FBS, 1% BSA (1 hour) and incubated with primary antibodies overnight at room temperature (RT) for organoids or two nights at 4°C for explants. After removal of the primary antibody solution, dilutions of secondary antibodies and DAPI in PBS were added to organoids for 6h at RT and overnight at RT for explants. Explants were mounted on a microscope slide using Aqua-Polymount. Organoids were stored in a 1:1 ratio of PBS and glycerol solution. Explants and organoids were imaged by confocal microscopy. A list of all the antibodies used is provided in Table S1.

### Relation between cross section area and number of cells composing organoids

Measure of the roundness of organoids was performed on mask resulting from organoids segmentation, using the formula: 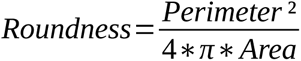

z-stack images of organoids were acquired at different stages of growth and the number of cells composing each organoid was manually counted. The relationship between the area of the central cross section of a sphere and its total volume follows the form:

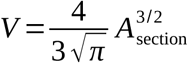

With *V* the volume of the sphere and *A_Section_* the area of the central cross section. Since the structure formed by organoids is very spherical, we fitted the data obtained with an equation of the same form and obtained the following equation: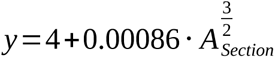

### Immunostaining-based quantitative image analysis

Segmentation of the organoids was performed on the DAPI channel. The channel was first blurred using a Gaussian filter and then segmented based on Otsu thresholding. The masks were manually corrected if necessary. Embryonic explants were manually segmented.

Nuclei were segmented in 2D using a pre-trained Stardist model (“2D_versatile_fluo”). We generated a map of the distance between each pixel within an organoid mask and the edge of the mask using the “findContours”, “drawContours” and “distanceTransform” functions of the OpenCV library. We calculated the center of mass of each nucleus mask and used the distance map to classify each cell as ‘external’ (distance to edge < 10 µm for embryonic tissues) or ‘internal’ (distance to edge > 10 µm for embryonic tissues).

#### p63 intensities quantification

After nuclear segmentation based on the DAPI channels, the resulting masks were used to calculate the average signal within each mask for normalized Hes1-emGFP or p63 channels. Statistical analysis was performed using the non-parametric Wilcoxon under R.

#### ECAD, ITGA6 and NMII-A quantification

After segmentation of the organoids’ global structure and of single nuclei as previously described, the ECAD, ITGA6 and NMII-A were normalized and each pixel assigned to a nucleus based on their distance to neighbouring nuclei calculated with a Voronoi diagram based on Euclidian distances. Statistical analysis was performed using the non-parametric Wilcoxon under R.

#### Profile of immunostaining intensity along distance to the BM in explants

Fluorescence intensity profiles were acquired from three independent biological replicates per tissue. Each profile consisted of measurements of signal intensity as a function of the normalized distance from the tissue edge toward the interior. Profiles were first smoothed individually using a centered moving average over a 10-point sliding window. Missing values generated at the beginning or end of the windowed series were replaced by the nearest available non-missing value to preserve edge data. Smoothed profiles were then normalized within each replicate by dividing all intensity values by the 95th percentile of that replicate’s smoothed intensity distribution. Because the distance sampling points were not identical across replicates, each profile was linearly interpolated to a common, tissue-specific grid of 200 equally spaced distance values spanning the full observed range. Mean normalized intensity and the standard error of the mean (SE) were then computed for each grid point across replicates. These mean ± SE values were used to generate tissue-specific profiles, to which linear regression was applied to estimate the relationship between normalized intensity and distance, reporting the coefficient of determination (R²) and p-value for the slope.

#### Quantification of Δ intensity under DAPT treatment

To quantify the effect of DAPT treatment on epithelial intensity profiles, we computed the change in mean fluorescence intensity between proximal and distal regions of the tissue. For each replicate, fluorescence intensity values were first extracted along a normalized distance axis (0 = tissue-stroma boundary, 50 = most distal region) from segmented epithelial compartments. Distances were binned into 1 µm intervals, and the mean intensity was calculated within each bin.

For each replicate, the proximal intensity was defined as the mean intensity in the 0-10 µm region, and the distal intensity was defined as the mean intensity in the 40-50 µm region. The Δ intensity was calculated as the absolute value of the difference between the two means for each biological replicate in both DAPT-treated and control conditions. Statistical differences between conditions were assessed using a two-sided Mann-Whitney U test.

### Organoid time-lapse image analysis

#### Tracking performance evaluation

Tracking performance was quantitatively evaluated against manually curated ground-truth lineages for five independent organoid movies. For each dataset, predicted tracks were compared to ground truth annotations using the built-in comparison framework of OrganoidTracker. Position-level detection accuracy was assessed using precision, recall and F1-score, based on a maximum spatial matching distance of 5 µm. Link accuracy was quantified analogously by comparing temporal associations between matched detections. Division detection performance was evaluated from lineage comparison categories, computing true positives (correctly detected divisions), false negatives (missed divisions), and false positives (spurious divisions), from which division recall and precision were derived. All metrics were computed over the full duration of each movie.

#### Fate classification from Hes1-emGFP reporter signal

To assign a final fate to each lineage, *Hes1-emGFP* intensity was normalized within each movie and time point by dividing the per-cell signal by the population mean at that movie/time point combination. For each lineage, a late Hes1 score was defined as the median normalized signal across the final 10 time points. Lineages were assigned to one of three fate classes, Hes1^high^ (luminal), Hes1^low^ (basal), or Hes1^int^ (stem-like/intermediate), using fixed thresholds on this late score (T∼high∼ = 1.0225, T∼low∼ = 0.9650). For analyses requiring an ordered variable, classes were encoded as integer ranks (Hes1^low^ = 0, Hes1^int^ = 1, Hes1^high^ = 2).

#### Lineage-level niche residency metric in pre-anchor windows

To quantify pre-anchor niche residency, we summarized each lineage’s distance-to-boundary trajectory within a specified pre-anchor window [t_0_, t_1_[ ending at the anchor (e.g. [−50,0[ minutes). For each lineage, all distance measurements falling within the window were collected and sorted by aligned time. We computed a time-averaged distance-to-edge metric:

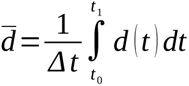

where *d* ( *t* ) is the absolute distance to the boundary and *Δ t* is the time span covered by that lineage within the window. The integral was approximated using the trapezoidal rule. Lineages were retained only if they met minimal coverage criteria (minimum number of samples and minimum covered duration within the window; thresholds reported in code).

#### Predictive association between pre-anchor niche residency and fate

To test whether pre-anchor niche residency predicts final Hes1 class, we evaluated the association between the lineage-level residency and the ordinal fate variable (0/1/2). We quantified monotonic association using Spearman rank correlation. To account for nesting of lineages within movies (organoid replicates), statistical significance was assessed using a movie-stratified permutation test: within each permutation, fate labels were randomly permuted within each movie while preserving per-movie class composition; Spearman correlation was then recomputed on the pooled permuted dataset. The empirical two-sided p-value was calculated as the fraction of permutations producing an absolute correlation greater than or equal to the observed value (with a +1 pseudocount).

#### Identification of the pre-commitment window

To identify which pre-symmetry-breaking time window carried the most fate-predictive positional information, we scanned windows of varying durations all ending at t = 0 (start times t₀ ∈ {−300, −250, −200, −150, −100, −75, −60, −50, −40, −30, −20, −15, −10, −5} minutes). For each window, the monotonic association between dh and the ordinal fate variable was quantified using Spearman rank correlation ρ. To account for the nesting of lineages within organoids, statistical significance was assessed using a within-organoid permutation test in which fate labels were permuted independently within each organoid across 20,000 permutations; the empirical two-sided p-value was the fraction of permutations producing |ρ| ≥ |ρ∼observed∼|. The window yielding the highest | ρ| among windows containing at least 30 lineages was selected as the pre-commitment window ([−50, 0) minutes; ρ = 0.427, permutation p = 5×10^−5^). Effect size was additionally quantified using an ordered logistic regression model, reporting the odds ratio per +1 µm increase in d with 95% confidence intervals.

#### Predictive accuracy of niche residency

To provide an intuitive measure of discriminative power between the two committed fate extremes, a standard ROC analysis was performed. The ROC curve plots the true positive rate against the false positive rate as the residency threshold for classifying a cell as luminal is varied across all possible values. The AUC summarizes this curve as a single number: AUC = 0.5 corresponds to chance discrimination, and AUC = 1.0 to perfect separation. A 95% bootstrap confidence interval was obtained from 2,000 resamples. As a supplementary analysis, one-vs-rest AUC was computed for each fate class independently to assess whether residency selectively predicts each class against the remaining two.

### Profile of immunostaining intensity along distance to the basement membrane in organoids

#### Depth profiles

The organoid contour was defined as the basement membrane, and the distance of each tissue pixel from this boundary was calculated using a Euclidean distance transform. Fluorescence intensity was measured within the tissue mask after subtraction of the median background intensity measured outside the organoids. Profiles from 0-20 µm from the basement membrane were divided into 0.5 µm bins. Mean fluorescence intensity was calculated for each bin, and bins containing fewer than 10 pixels were excluded.

#### Compartments and normalization

Two compartments were defined: external (0-5 µm from the basement membrane) and internal (5-20 µm). Mean intensity within each compartment was calculated by weighting each depth bin by its number of pixels. For each marker and comparison, intensities were normalized to the mean control intensity measured over 0-20 µm, such that the control mean equals 1 and treated samples are expressed as fold change. The acquisition was used as the unit of replication. Statistical comparisons were performed using a linear mixed-effects model: intensity ∼ condition + (1 | acquisition) with acquisition included as a random effect.

### Embryonic time-lapse image analysis

To remove spectral crosstalk, 3 channel images were spectrally unmixed into 3 images corresponding to the contribution from each of the 3 fluorophore species present in the sample. To this aim, reference spectra of the utilized fluorophores were determined from calibration images acquired on positive samples for only one fluorophore.

To compensate for axial (Z-axis) signal attenuation in confocal image stacks, we applied an automated correction procedure using the DAPI signal as a reference. 4D images (3D + time) containing DAPI and Hes1 channels were loaded together with a binary nuclear mask (generated by Stardist), which was rescaled to match the raw image voxel size. For each time point, the mean DAPI intensity was measured within the mask at each Z-slice, and a slice-specific correction factor was computed by normalizing to the mean intensity across slices.

## Supplementary Information

**Table S1:** Antibodies.

| Antibody | Source | Reference | Dilution |
|---|---|---|---|
| Rabbit anti-K5 | Covance | Cat# PRB-160P-100; RRID: AB_291581 | 1:300 |
| Rat anti-K8 | Developmental Studies Hybridoma Bank, University of Iowa | Cat# TROMA-I: RRID: AB_531826 | 1:300 |
| Mouse anti-p63 | Abcam | Cat# ab735; RRID: AB_305870 | 1:200 |
| Rabbit anti-Ki67 | Thermo Fisher Scientific | Cat# MA5-14520; RRID: AB_10979488 | 1:300 |
| Rabbit anti-HES1 | Cell Signalling technology | #11988 | 1:100 |
| Rabbit anti-NMII-A | Biolegend | Cat# 909801; RRID: AB_2734686 | 1:300 |
| Rabbit anti-LAMA332 | Kind gift of M. Aumalley, University of Cologne, Germany | Serum | 1:300 |
| Rabbit anti-NMII-B | Biolegend | Cat# 909802 | 1:300 |
| Mouse anti-Phospho-MLC/MYL9 | Invitrogen | Cat# MA5-15163; RRID: AB_10979087 | 1:150 |
| Rabbit anti-ECAD | Cell Signalling technology | Cat# 3195; RRID: RRID: AB_2291471 | 1:300 |
| Mouse anti-ECAD | BD Transduction Laboratories | Cat# 610182; RRID: RRID: AB_397581 | 1:200 |
| Rat anti-ITGA6 | BD Transduction Laboratories | Cat# 555734; RRID: RRID: AB_ 2296273 | 1:400 |
| Human anti-GFP | Recombinant Protein Platform, Institut Curie, Paris | Recombinant anti GFP antibody | 1:200 |
| Rabbit anti-YAP | Cell Signalling Technology | Cat# 14074; RRID: AB_2650491 | 1:300 |
| Goat anti-rabbit AlexaFluor-coupled to different fluorochromes (Cy3, Cy5, A488, A405) | Invitrogen-Thermo Fisher Scientific | Cat# A10520; RRID: AB_2534029 Cat#A10523; RRID: AB_2534032, Cat# A-11034; RRID: AB_2576217 Cat# A-31556; RRID: AB_221605 | 1:500 |
| Goat anti-rat AlexaFluor-coupled to different fluorochromes (Cy3, Cy5, A488) | Invitrogen-Thermo Fisher Scientific | Cat# A10522; RRID: AB_2534031, Cat#A10525; RRID: AB_2534034, Cat# A-11006; RRID: AB_2534074 | 1:500 |
| Goat anti-mouse AlexaFluor-coupled to different fluorochromes (Cy3, Cy5, A488) | Invitrogen-Thermo Fisher Scientific | Cat# A10521; RRID: AB_2534030, Cat# A10524; RRID: AB_2534033 Cat# A- 11001; RRID: AB_2534069 | 1:500 |
| EpCAM PE/Cy7 | Biolegend | Cat# 118216; RRID: AB_1236471 | 1:100 |
| CD49f APC/Cy7 | Biolegend | Cat# 313628; RRID: AB_2616784 | 1:100 |
| Pdpn APC/Cy7 | Biolegend | Cat# 127417; RRID: AB_2629803 | 1:100 |

**Table S2.** Reagents and concentration.

| Reagent | Source | Reference | Concentration | Organoid medium |
|---|---|---|---|---|
| DMEM/F12 | Gibco | 21331020 | 1x | MG, LG, SG, Pro |
| Glutamax | Thermo Fisher Scientific | 35050038 | 1x | MG, LG, SG, Pro |
| HEPES (1M) | Gibco | 15630080 | 1x | MG, LG, SG, Pro |
| B27 supplement | Thermo Fisher Scientific | 12587-010 | 2x | MG, LG, SG, Pro |
| N2 supplement | Thermo Fisher Scientific | 17502048 | 1x | MG |
| Penicillin/Streptomycin | Thermo Fisher Scientific | 15140122 | 2% | MG, LG, SG, Pro |
| FGF STAB | Enantis | FGF2STAB01000 | 50ng/mL | MG, LG, SG, Pro |
| mNoggin | PeproTech | 250-38 | 100 ng/mL | MG, LG, SG, Pro |
| mRspo1 | PeproTech | 3474-RS | 100 ng/mL | MG, LG, SG, Pro |
| N-acetylcysteine | Medchemexpress | HY-B0215 | 1.25 mM | SG, LG, Pro |
| Neuregulin | R&D Systems | 5898-NR-050 | 100ng/mL | MG |
| Wnt3a | R&D Systems | 1324-WNP-010/CF | 5 ng/mL | LG, SG |
| mEGF | Thermo Fisher Scientific | 315-09 | 2 ng/mL | LG, SG, Pro |
| A83-01 | TOCRIS | 2939 | 10 µM | Pro |
| Dihydro-testosterone (DHT) | Sigma-Aldrich | A8380 | 1 nM | Pro |
| Y-27632 | Medchemexpress | HY-10071 | 10 µM, 20/100 for perturbation of Myosin | MG, LG, SG, Pro |
| Tamoxifen | MP Biomedicals | 156738 | 1mg/10g of mouse |  |
| Cell Recovery Solution | Corning | 354253 | 1x |  |
| CO2 independent medium | Gibco | 18045054 | 1x |  |
| Fetal bovine serum (FBS) | Thermo Fisher Scientific | 10500064 | 10% |  |
| Insulin Transferrin Selenium (ITS) | Gibco | 41400045 | 2% |  |
| Matrigel | Corning | 354234 | 5-10mg/mL |  |
| AggreWell™400 | STEMCELL technologies | #34415 |  |  |

**Figure S1:**
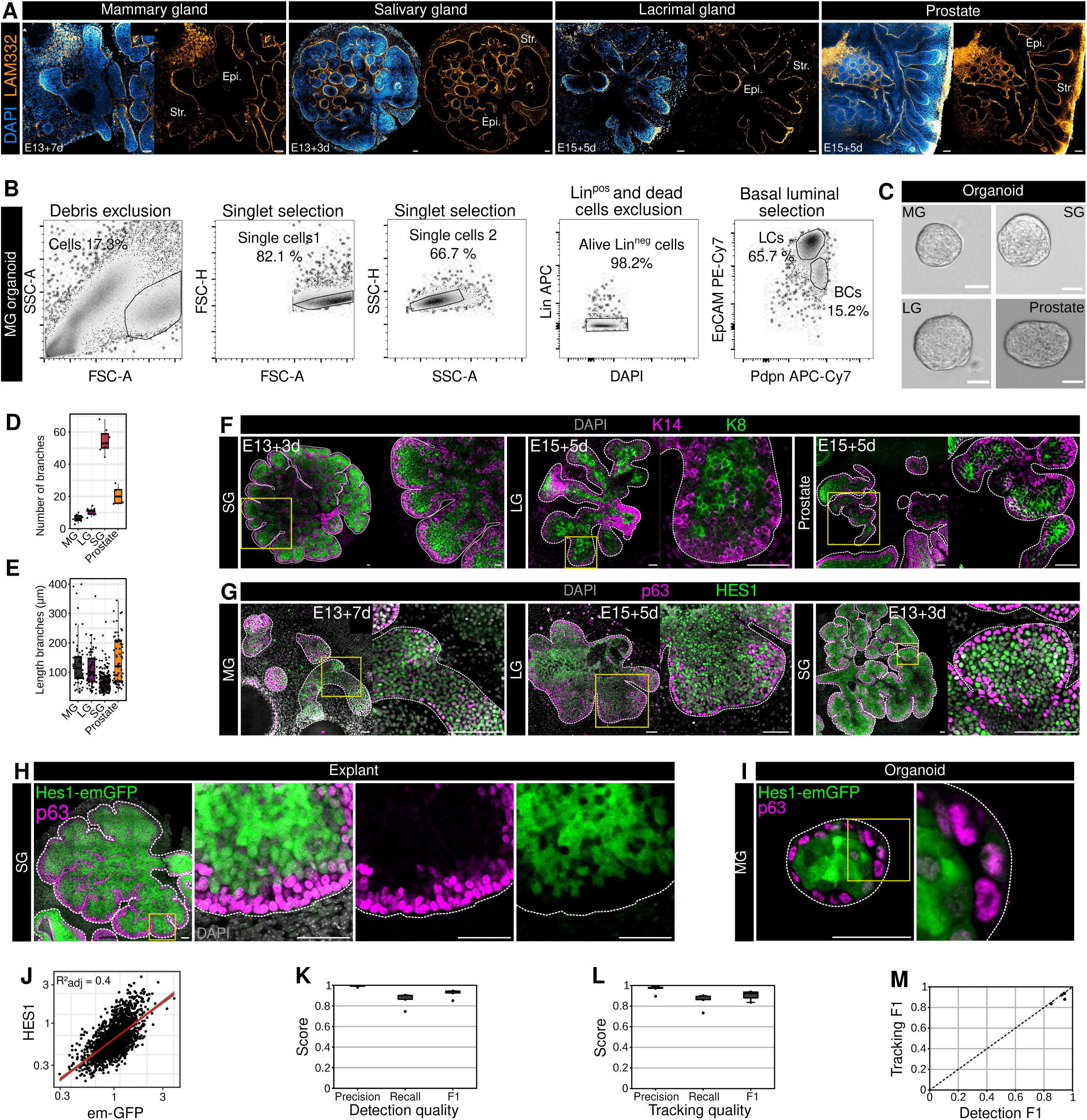
*Ex-vivo* explant and organoid models recapitulate glandular epithelial organization and enable quantitative cell tracking. **A:** Confocal imaging of MG, SG, LG, and prostate embryonic explants after the indicated periods of *ex vivo* culture. Epi: epithelium, Str: stroma. **B:** Representative FACS gating strategy used to isolate mammary organoid epithelial populations. Debris and doublets were sequentially excluded, followed by selection of live lineage-negative (Lin^neg^) cells and separation of basal cells (BCs) and luminal cells (LCs) based on EpCAM and podoplanin (Pdpn) expression. **C:** Representative brightfield images of organoids generated from MG, SG, LG, and prostate epithelial cells. **D:** Quantification of the number of branches formed by MG, LG, SG, and prostate embryonic explants during *ex vivo* culture. **E:** Quantification of branch length in MG, LG, SG, and prostate embryonic explants. **F:** Confocal imaging of K14 (magenta) and KS (green) in SG, LG, and prostate embryonic explants after the indicated periods of *ex vivo* culture. **G:** Confocal imaging of p63 (magenta) and HES1 (green) in MG, LG, and SG embryonic explants after the indicated periods of *ex vivo* culture. **H:** Confocal imaging of a Hes1-emGFP SG embryonic explant stained for p63, illustrating the spatial segregation of HES1-high and p63-positive cells *ex vivo*. **I:** Confocal imaging of a Hes1-emGFP mammary organoid stained for p63, illustrating the corresponding spatial organization *in vitro*. **J:** Relation between HES1 and Hes1-emGFP intensities in nucleus of MG organoids cells. Spearman’s rank correlation. **K:** Performance of the automated cell-detection pipeline, quantified by precision, recall, and F1 score. **L:** Performance of the automated cell-tracking pipeline, quantified by precision, recall, and F1 score. **M:** Relationship between detection F1 score and tracking F1 score across analyzed movies. Scale bars: 50 μm. Dotted lines in F, G and H, delineate the epithelium limit.

**Figure S2:**
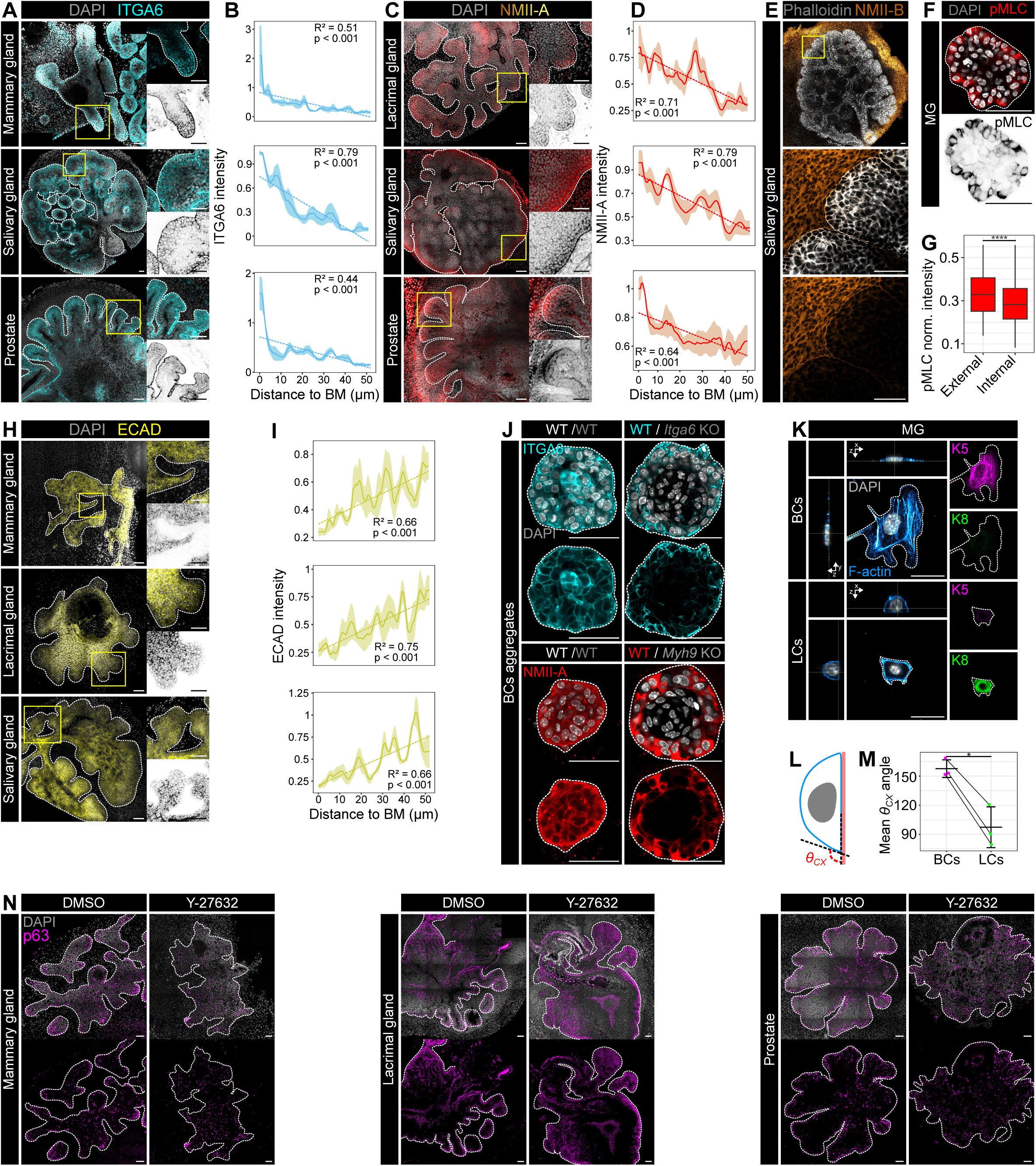
Adhesion-contractility components display compartment-specific organization across glandular epithelia. **A:** Confocal imaging of ITGA6 in MG, SG, and prostate embryonic explants. **B:** Mean ITGA6 intensity as a function of distance from the BM in the corresponding tissues. **C:** Confocal imaging of NMII-A in lacrimal gland, salivary gland and prostate embryonic explants. Insets show higher-magnification views. **D:** Mean NMII-A intensity as a function of distance from the BM in the corresponding tissues. **E:** Representative confocal imaging of F-actin, visualized with phalloidin, and NMII-B in salivary explants, illustrating their spatial organization across the epithelium. **F:** Confocal imaging of phosphorylated myosin light chain (pMLC) in mammary organoids. **G:** Quantification of normalized pMLC intensity in the external and internal epithelial compartments. **H:** Confocal imaging of ECAD in mammary, lacrimal, and salivary gland embryonic explants. **I:** Mean ECAD intensity as a function of distance from the BM in the corresponding tissues. **J:** Confocal imaging of basal-cell aggregates composed of WT cells alone or mixtures of WT cells with ITGA6-KO or NMII-A-deficient cells, stained for the indicated adhesion-contractility component. **K:** Representative confocal imaging of FACS-sorted mammary BCs and LCs plated on Matrigel coated glass slides. F-actin is shown together with the basal marker K5 or luminal marker KS. Orthogonal views illustrate cell morphology and spreading on the substrate. **L:** Schematic representation of the cell–substrate contact angle *θ*_CX_ used to quantify cell spreading. **M:** Mean cell-substrate contact angle of basal and luminal mammary epithelial cells. Lines connect measurements obtained from matched biological replicates. **N:** Confocal imaging of p63 in mammary gland, lacrimal gland, and prostate embryonic explants treated with DMSO or 20 μM of the ROCK inhibitor Y-27632. DAPI is shown in grey and p63 in magenta. Scale bars, 50 μm. Dotted lines in A, C, E, F, H, J and N delineate the epithelium limit.

**Figure S3:**
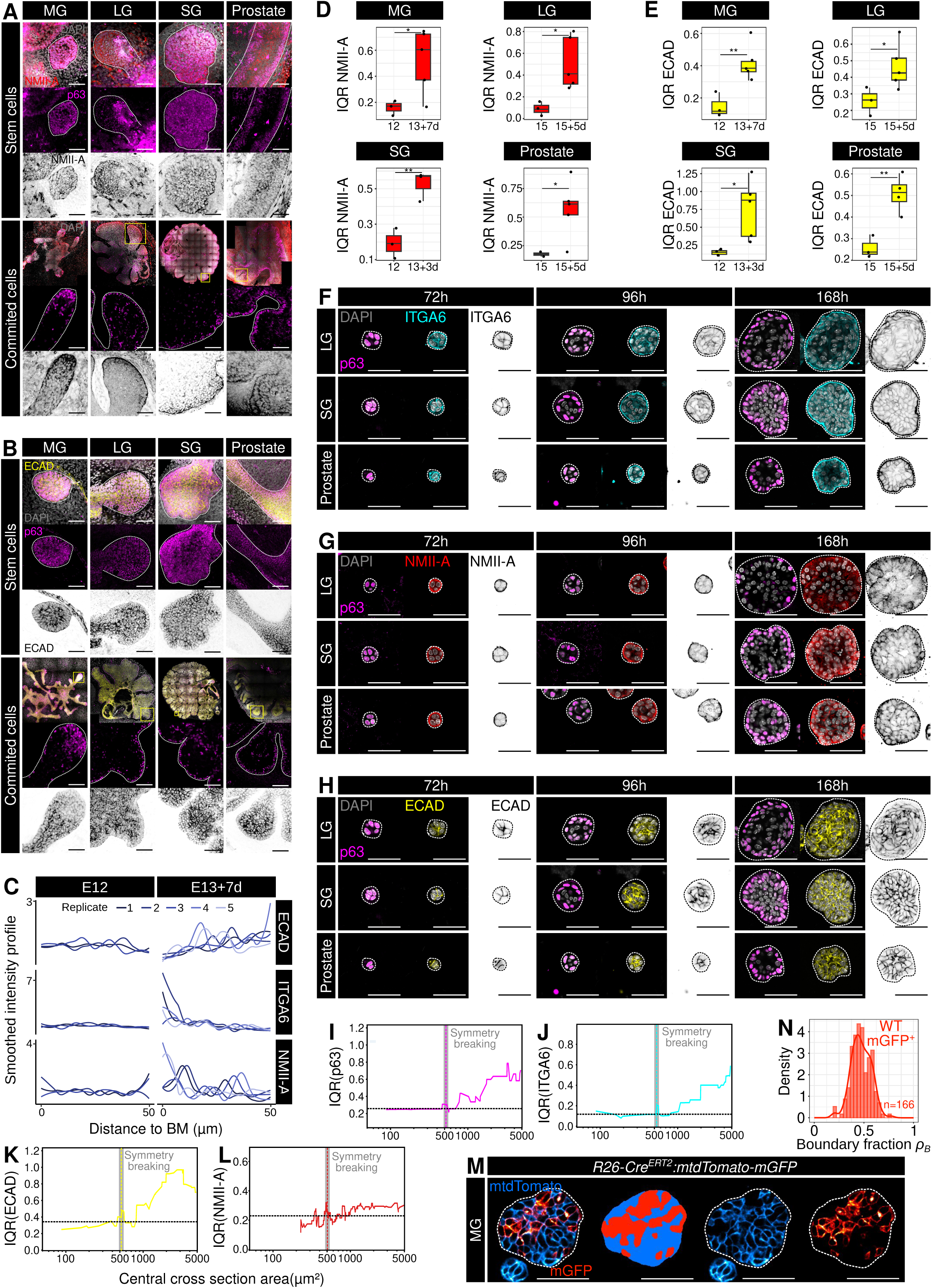
Adhesion-contractility heterogeneity progressively emerges during epithelial lineage commitment. **A:** Confocal imaging of NMII-A in MG, LG, SG, and prostate epithelia at early, stem-like and later, lineage-committed stages of development. **B:** Confocal imaging of ECAD in the same tissues and developmental stages. **C:** Smoothed intensity profiles of ECAD, ITGA6, and NMII-A as a function of distance from the BM at early and later stages of embryonic mammary gland development. Individual lines represent independent biological replicates. **D:** Mean IQR of cellular NMII-A intensity at early and later developmental stages in MG, LG, SG, and prostate epithelia. **E:** Mean IQR of cellular ECAD intensity at early and later developmental stages in the same tissues. **F-H:** Confocal imaging of ITGA6, NMII-A and ECAD in LG-, SG-, and prostate-derived organoids after 72, 96, and 16S h of culture. **I-L:** Interquartile range of p63 (**I**), ITGA6 (**J**), ECAD (**K**), and NMII-A (**L**) intensity as a function of organoid central cross-sectional area. Vertical lines indicate the onset of S-16 transition. **M:** Representative confocal image of an organoid derived from *R26-Cre^ERT^*^2^*;mTdTomatolmGFP* cells, showing the spatial distribution of unrecombined tdTomato-positive and recombined mGFP-positive cells. The corresponding segmented populations are shown in the adjacent panel. **N:** Probability-density distribution of the boundary fraction p_B_ occupied by WT mGFP-positive cells in mosaic mammary organoids. Scale bars, 50 μm. Dotted lines in A, B, F-H and M delineate the epithelium limit.

**Figure S4:**
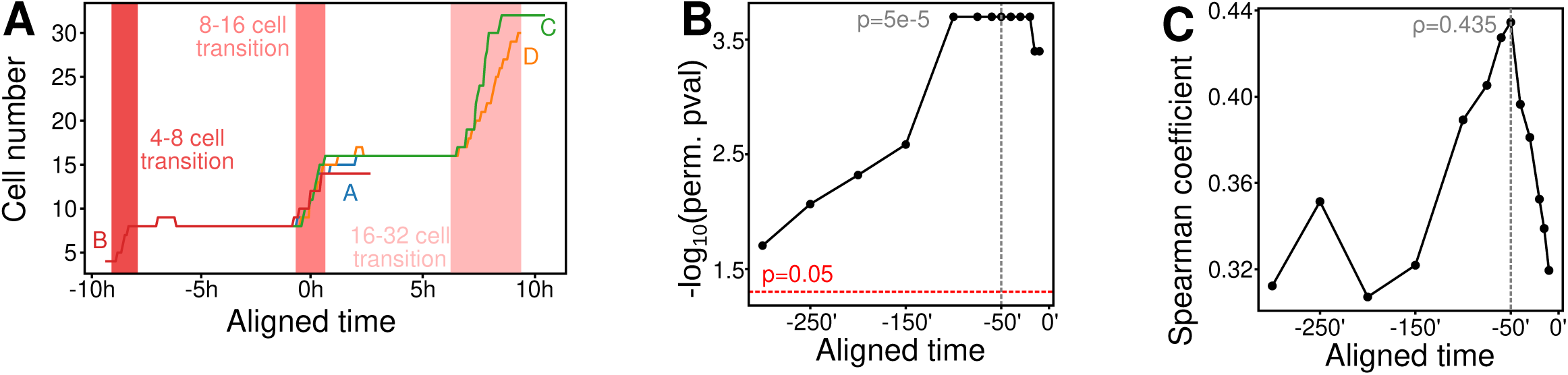
Positional information emerges before lineage stabilization during organoid growth. **A:** Representative cell-number trajectories from individual organoids aligned to the onset of HES1 symmetry breaking. Shaded regions indicate the 4-S-, S-16-, and 16-32-cell transitions. **B:** Temporal evolution of the statistical association between pre-commitment cell position and subsequent HES1 state. Significance was assessed by permutation testing and is shown as −(P_perm_) . The red dashed line indicates P = *0*.*05*, and the vertical dashed line indicates 50 min before HES1 symmetry breaking. **C:** Spearman correlation coefficient between cell position and subsequent HES1 state as a function of time before symmetry breaking. The vertical dashed line indicates 50 min before HES1 symmetry breaking. The Spearman correlation at this time point is indicated.

**Figure S5:**
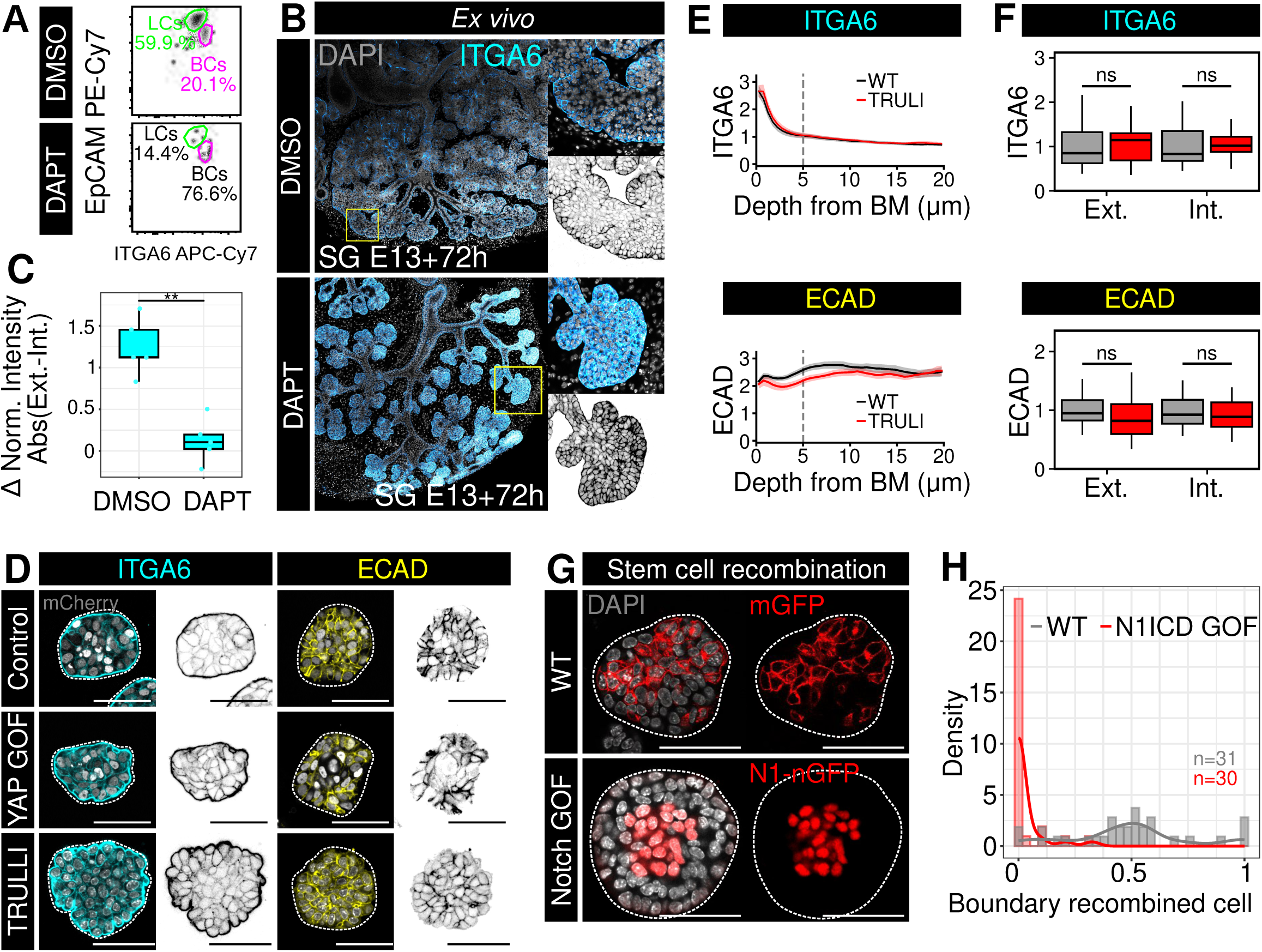
Notch and YAP pathway differentially regulate epithelial adhesion and cell positioning. **A:** Representative flow-cytometry analysis of EpCAM and ITGA6 expression following treatment with DMSO or the Notch inhibitor DAPT. Gates indicate LCs and BCs, with the percentage of cells in each population indicated. **B:** Confocal imaging of ITGA6 in salivary gland explants (E13+3d) cultured for 3 days *ex vivo* under DMSO or DAPT treatment. **C:** Quantification of the absolute difference in normalized ITGA6 intensity between the external and internal epithelial compartments following DMSO or DAPT treatment. **D:** Confocal imaging of ITGA6 and ECAD in control organoids and following genetic or pharmacological YAP activation, using YAP gain-of-function (YAP^S127A^) or TRULI, respectively. **E:** Mean ITGA6 and ECAD intensity profiles as a function of distance from the BM in WT and TRULI-treated organoids. The vertical dashed line indicates the boundary between the external and internal compartments. **F:** Quantification of ITGA6 and ECAD levels in the external (Ext.) and internal (Int.) compartments of WT and TRULI-treated organoids. **G:** Representative confocal imaging of control and Notch gain-of-function organoids following recombination in early phase of organoid growth (OHT during the initial 4Sh). Recombined control cells are identified by mGFP, whereas Notch-activated cells are identified by N1-nGFP. **H:** Probability-density distribution of the boundary localization of recombined cells in control (WT) and N1ICD gain-of-function organoids. Scale bars, 50 μm. Dotted lines in B, D and G delineate the epithelium limit.

**Supplementary Videos 1-4: Branching morphogenesis of embryonic glandular epithelia ex vivo. Explant cultures recapitulating embryonic epithelial branching within native stroma *ex vivo*.**

- **Supplementary Video 1:** Mammary gland *ex vivo* branching morphogenesis. Live imaging of embryonic mammary gland explants. T0 corresponds to embryonic day E13 + 5 days in culture (E13+5d). Scale bar, 50 μm.
- **Supplementary Video 2:** Lacrimal gland *ex vivo* branching morphogenesis. Live imaging of embryonic lacrimal gland explants. T0 corresponds to E15+3d. Scale bar, 50 μm.
- **Supplementary Video 3:** Salivary gland *ex vivo* branching morphogenesis. Live imaging of embryonic salivary gland explants. T0 corresponds to E13+2d. Scale bar, 50 μm.
- **Supplementary Video 4:** Prostate *ex vivo* branching morphogenesis. Live imaging of embryonic prostate explants. T0 corresponds to E15+3d. Scale bar, 50 μm.

**Supplementary Video 5:** Live imaging of Hes1 dynamics during salivary gland branching morphogenesis. Time-lapse movie of *NLS-mKate2:Hes1-emGFP* salivary gland explants undergoing *ex vivo* branching morphogenesis. T0 corresponds to E13+2d. Images are acquired every 5 minutes. Scale bar, 50 μm.

**Supplementary Video 6:** Live imaging of *NLS-mKate2:Hes1-emGFP* mammary organoids after lineage commitment. Time-lapse movie of mammary organoids following lineage commitment. T0 corresponds to day 5 of culture (5d). Images are acquired every 5 minutes. Scale bar, 25 μm.

**Supplementary Video 7:** Inter-compartment cell exchanges during salivary gland branching morphogenesis in a *NLS-mKate2:Hes1-emGFP* mouse. High-magnification tracking showing cell exchanges between basal and luminal compartments during salivary gland explant morphogenesis (yellow arrows). Images are acquired every 5 minutes. Scale bar, 25 μm.

**Supplementary Video 8:** Live imaging of *NLS-mKate2:Hes1-emGFP* mammary organoids after lineage commitment. Identification of In-Out-In and Out-In-Out movements (yellow arrows). Images are acquired every 5 minutes. Scale bar, 25 μm.

**Supplementary Video 9:** Live imaging of *NLS-mKate2:Hes1-emGFP* mammary organoids at the onset of symmetry breaking. Time-lapse movie tracking mammary organoid development during early symmetry breaking. T0 corresponds to day 2 of culture (2d). Yellow arrows indicate the onset of symmetry breaking. Scale bar, 25 μm.

**Supplementary Video 10:** Live imaging of *NLS-mKate2* mammary organoids under DMSO or DAPT treatment. Time-lapse movie comparing control (DMSO) and Notch-inhibited (DAPT) *NLS-mKate2* mammary organoids. Treatments were applied at T0 (day 5 of culture) and maintained for the full duration of imaging. Scale bar, 25 μm.

**Supplementary Video 11:** Live imaging of *NLS-mKate2* mammary organoids under DMSO or TRULI treatment. Time-lapse movie comparing control (DMSO) and TRULI-treated *NLS-mKate2* mammary organoids. Treatments were applied at T0 (day 5 of culture) and maintained for the full duration of imaging. Scale bar, 25 μm.

**Supplementary Video 12**: Live imaging of inducible YAP-S127A expression in mammary organoids. Time-lapse movie of *R26-rtTA;tetO-YAP-S127A;tetO-mCherry* and control (*R26-rtTA;tetO-mCherry*) mammary organoids under doxycycline treatment. Doxycycline was added at T0 (day 5 of culture) and maintained throughout the full culture period. Scale bar, 50 μm.

## Notes

### Competing Interest Statement

The authors have declared no competing interest.

